# Biomechanics of aging and osteoarthritic human knee ligaments

**DOI:** 10.1101/2021.11.21.469435

**Authors:** Abby E Peters, Brendan Geraghty, Karl T Bates, Riaz Akhtar, Rosti Readioff, Eithne Comerford

**Affiliations:** Department of Mechanical, Materials and Aerospace Engineering, School of Engineering, University of Liverpool, Liverpool, L69 3GH, United Kingdom; Institute of Life Course and Medical Sciences, University of Liverpool, Liverpool, L7 8TX, United Kingdom; Medical Research Council Versus Arthritis Centre for Integrated Research into Musculoskeletal Ageing (CIMA), University of Liverpool, Liverpool, United Kingdom; Institute of Medical and Biological Engineering, School of Mechanical Engineering, Faculty of Engineering, University of Leeds, Leeds, UK; School of Veterinary Science, University of Liverpool, Neston, CH64 7TE, United Kingdom

## Abstract

**Background:** Ligaments work to stabilize the human knee joint and prevent excessive movement. Whilst ligaments are known to decline in structure and function with aging, there has been no systematic effort to study changes in gross mechanical properties in the four major human knee ligaments due to osteoarthritis (OA). This study aims to collate material properties for the anterior (ACL) and posterior (PCL) cruciate ligaments, medial (MCL) and lateral (LCL) collateral ligaments. Our cadaveric samples come from a diverse demographic from which the effects of aging and OA on bone and cartilage material properties have already been quantified. Therefore, by combining our previous bone and cartilage data with the new ligament data from this study we are facilitating subject-specific whole-joint modelling studies.

**Methods:** The demographics of the collected cadaveric knee joints were diverse with age range between 31 to 88 years old, and OA International Cartilage Repair Society grade 0 to 4. Twelve cadaveric human knee joints were dissected, and bone-ligament-bone specimens were extracted for mechanical loading to failure. Ligament material properties were determined from the load-extension curves, namely: linear and ultimate (failure) stress and strain, secant modulus, tangent modulus, and stiffness.

**Results:** There were significant negative correlations between age and ACL linear force (*p=0.01*), stress (*p=0.03*) and extension (*p=0.05*), ACL failure force (*p=0.02*), stress (*p=0.02*) and extension (*p=0.02*), PCL secant (*p=0.02*) and tangent (*p=0.02*) modulus, and LCL stiffness (*p=0.05).* Significant negative correlations were also found between OA grades and ACL linear force (*p=0.05*), stress (*p=0.02*), extension (*p=0.01*) and strain (*p=0.03*), and LCL failure stress (*p=0.05*). However, changes in age or OA grade did not show a statistically significant correlation with the MCL tensile parameters. Trends showed that almost all the tensile parameters of the ACL and PCLs decreased with increasing age and progression of OA. Due to small sample size, the combined effect of age and presence of OA could not be statistically derived.

**Conclusions:** This research is the first to correlate changes in tensile properties of the four major human knee ligaments to aging and OA. The current ligament study when combined with our previous findings on bone and cartilage for the same twelve knee cadavers, supports conceptualization of OA as a whole-joint disease that impairs the integrity of many peri-articular tissues within the knee. The subject-specific data pool consisting of the material properties of the four major knee ligaments, subchondral and trabecular bones and articular cartilage will aid reconstruction and graft replacements and advance knee joint finite element models, whilst knowledge of aged or diseased mechanics may direct future therapeutic interventions.

## Introduction

The human knee joint is composed of both soft and hard tissues, forming a diarthrosis articulation between the femur and tibia, allowing predominantly flexion and extension in the sagittal plane (Nigg & Herzog, 2007). Primary human knee joint ligaments act as strain sensors, constraining degrees of freedom to provide stabilization and prevent excessive movement (Harner et al., 1995; Woo et al., 2006). Structurally, ligaments have direct and indirect insertions into the bone and periosteum (Woo et al., 2006) allowing fiber bundle variations to respond to different movements and resist loading during tension or rotation at the joint (Hansen, Masouros & Amis, 2006).

Tensile properties of the anterior cruciate ligament (ACL), posterior cruciate ligament (PCL), medial collateral ligament (MCL) and lateral collateral ligament (LCL) have been explored by numerous researchers (Noyes & Grood, 1976; Woo et al., 1991; Race & Amis, 1994; Robinson, Bull & Amis, 2005; Bonner et al., 2015; Smeets et al., 2017; Cho & Kwak, 2020; Patel et al., 2021), providing important information on their structural and mechanical properties. Data for all four ligaments exists across various sources in the literature; however, these are most often harvested and tested in isolation, obtaining just one ligament type from donors. To date, very few studies have explored all four ligaments from the same donor (donors were limited to healthy knee joints), with data suggesting higher stiffness and failure load in the cruciate ligaments compared to the collateral ligaments (Trent, Walker & Wolf, 1976; van Dommelen et al., 2005). Despite data existing for all four ligaments from varying specimens in previous studies, there is marked variability in reported values, likely due to variations in testing techniques and donor demographics, which currently makes it challenging to understand whole-joint function (Peters et al., 2018a).

The lack of consistent healthy baseline measurements means our understanding of how tensile properties of all four ligaments within the same knee joint change with aging or disease is presently unclear (Peters et al., 2018a). Structural and functional capabilities are known to decline with age in the ACL, in particular a decrease in ultimate failure load from older donors (67 to 90 years), when compared to donors between 40 to 50 years old, and younger donors (22 to 35 years) (Woo et al., 1991). This is also reflected at a cellular level in ligaments such that ACL histological parameters, including cartilage fiber disorganization and mucoid degeneration, showed an increase in tissue degeneration with age (Hasegawa et al., 2012). However, any differences in material properties in the PCL, MCL and LCL are yet to be systematically correlated with different age categories. Changes to integrity and tensile properties not only leave ligaments vulnerable to further injury but also affect the peri-articular tissues leading to muscle weakening through immobility, and whole-joint disruption including the development of osteoarthritis (OA) (Rousseau & Garnero, 2012; Manninen et al., 1996; Simon et al., 2015). In addition, our knowledge about the effect of OA on the tensile properties of the knee joint ligaments is limited, with current studies focusing primarily on histological analyses. There is evidence showing impaired integrity of the ACL and PCL during total knee replacements in the presence of OA and with age (Mullaji et al., 2008; Hasegawa et al., 2012).

It is valuable to link the previously observed data on histological and micro-scale morphological changes due to aging and OA in the knee joint ligaments, to changes in the tissue-level ligament mechanics, and then to the whole knee joint mechanics (Kumar, Manal & Rudolph, 2013; Adouni, Shirazi-Adl & Shirazi, 2012). Previously, we systematically investigated the effect of age and OA on the mechanical properties of bone and cartilage in human knee joints for the first time on the same donor (Peters et al., 2018b). Here, we have employed the same human cadavers to study the ligaments. This therefore will allow (i) the first assessment of changes in the mechanical behavior of ligaments due to aging and OA, (ii) ligament data to be combined with bone and cartilage trends from the same specimen to give a fuller picture of multi-tissue joint (whole-joint) changes with age and OA, and (iii) the subject-level data to be used in future development of subject-specific OA knee joint computer models. Thus, the aim of this study was to obtain data on tissue-level material characteristics of cadaveric human knee joint ligaments with a wide span of age and OA grades and correlate these to age and OA grade as univariate parameters. To fulfill the aim of this study, the following objectives were performed:

1. Harvesting the four major knee joint ligaments (ACL, PCL, LCL and MCL) as a bone-ligament-bone specimens, and measure undeformed geometrical parameters of the ligaments.
2. Application of physiologically relevant tensile loads on the ligaments and determining their mechanical responses.
3. Analysis of the tensile characteristics of the ligaments and tests of their correlations with age and OA.

## Materials & Methods

### Specimens

Fresh-frozen human cadaveric knee joints were sourced from Science Care (Phoenix, Arizona, United States) via Newcastle Surgical Training Centre (Newcastle upon Tyne, NE7 7DN, United Kingdom); therefore, the consents were obtained and held by Science Care. The knee cadavers were from humans aged 31 to 88 years (n = 12; 4 female and 8 male) (Table S2) as reported in our previous study (Peters et al., 2018b). Ethical permission for use of this human cadaveric material was sponsored by the University of Liverpool and granted by the National Research Ethics Service (15/NS/0053) who approved all protocols. All experiments were performed in accordance with relevant guidelines and regulations.

Cadaver limbs were initially frozen at −20°C and thawed at 3 to 5°C for 5 days prior to dissection. During dissection, cadavers were photographed and graded for OA using the International Cartilage Repair Society (ICRS) (Supplementary Materials (Table S1)) as reported in our previous study (Peters et al., 2018b). Four bone-ligament-bone specimens were harvested from each cadaver using a low-speed oscillating saw (deSoutter Medical, Bucks, UK) (Fig. 1A). Extracted specimens were then stored at −20°C before they were thawed for 24 hours at 3 to 5°C and submerged in phosphate buffered saline (PBS). Overall the specimens underwent two freeze-thaw cycles, which has previously been shown to have no effect on ligament and tendon material properties (Woo et al., 1986; Moon et al., 2006; Huang et al., 2011; Jung et al., 2011; Peters et al., 2017). Specimen numbers are consistent with those in Peters et al. (2018b), allowing matching of ligament properties presented here with previously reported cartilage and bone data from the same individuals (Supplementary Materials (Ligament Raw Data.xlsx)).

**Fig. 1.**
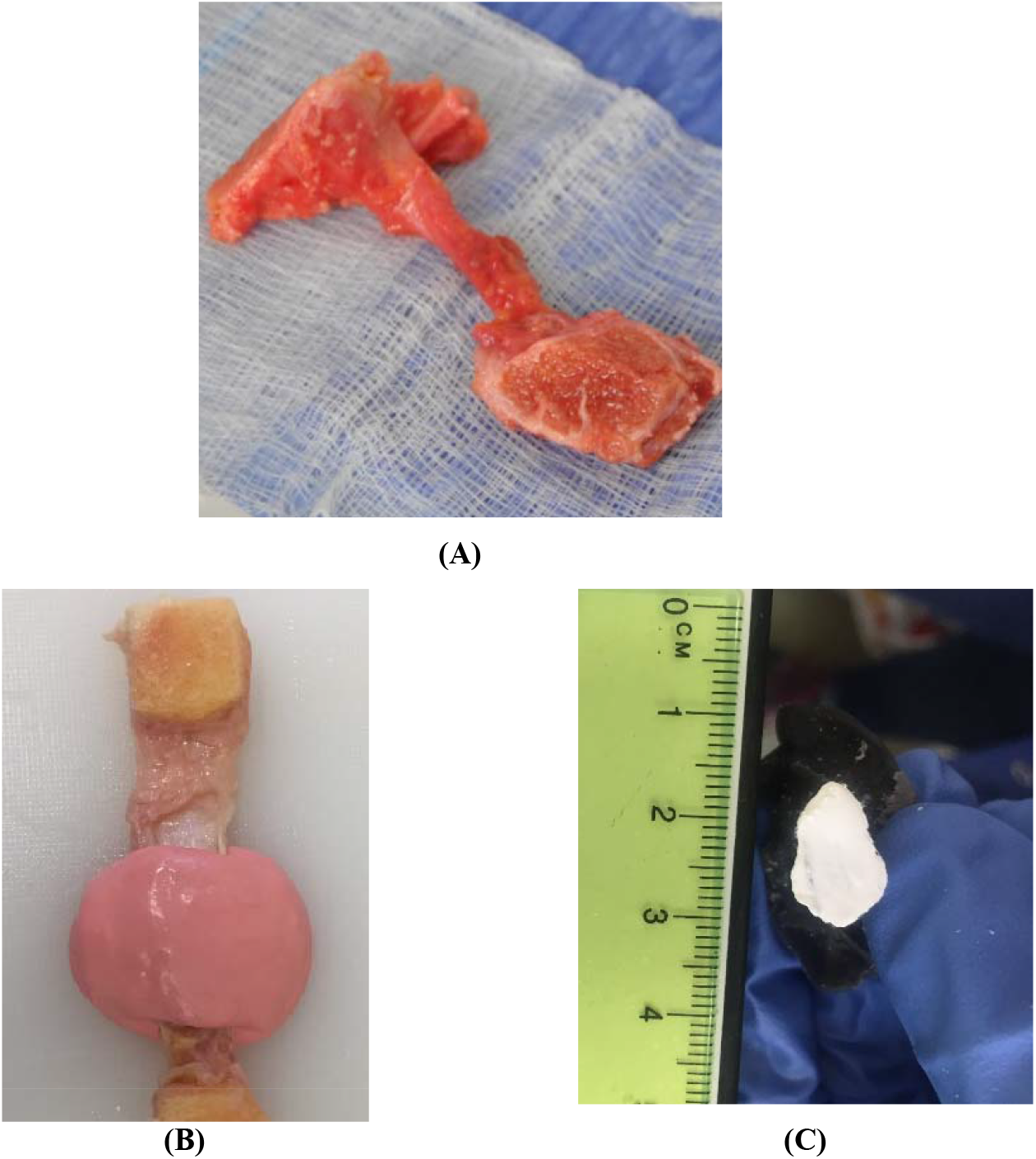
Bone-ligament-bone preparation and method for measuring the middle cross-sectional area of knee joint ligament specimens. (A) Bone-ligament-bone specimen. (B) Ligament encased in impression material. (C) Polymethyl-methacrylate cast of a ligament photographed for crosssectional area measurement.

### Cross-Sectional Area Measurements

The cross-sectional area of all of the ligament specimens were obtained using a previously described method (Goodship & Birch, 2005; Readioff et al., 2020b). In brief, using a fast-setting alginate impression paste (UnoDent, Essex, England) ligaments were encased in the material and left to set for two minutes (Fig. 1B). Once the impression material was set, a scalpel blade was used to slice the mold which was then filled with polymethyl-methacrylate (PMMA) (Teknovit 6091, Heraeus Kulzer GmbH, Wehrheim, Germany) to create a replica of the ligament structure. Once the PMMA was set, the mold was sliced transversely, and the resulting ends colored with permanent white marker pen (Fig. 1C). The cement mold ends were then photographed and digitally measured using ImageJ (Schneider, Rasband & Eliceiri, 2012) to obtain the cross-sectional area of the ligament.

### Specimen Preparation

Bone around the insertion and origin sites of each ligament on the femur and tibia were cut into a suitable shape using a hand saw (Fig. 1A). The bone ends of the specimens were potted into custom-made stainless-steel holders and screwed in place. PMMA was then poured into the holder and left to cure for 4 to 5 minutes. Specimens were then attached to the load cell and encased into a watertight custom-made chamber. The chamber was filled with PBS to control specimen hydration during testing (Fig. 2).

**Fig. 2.**
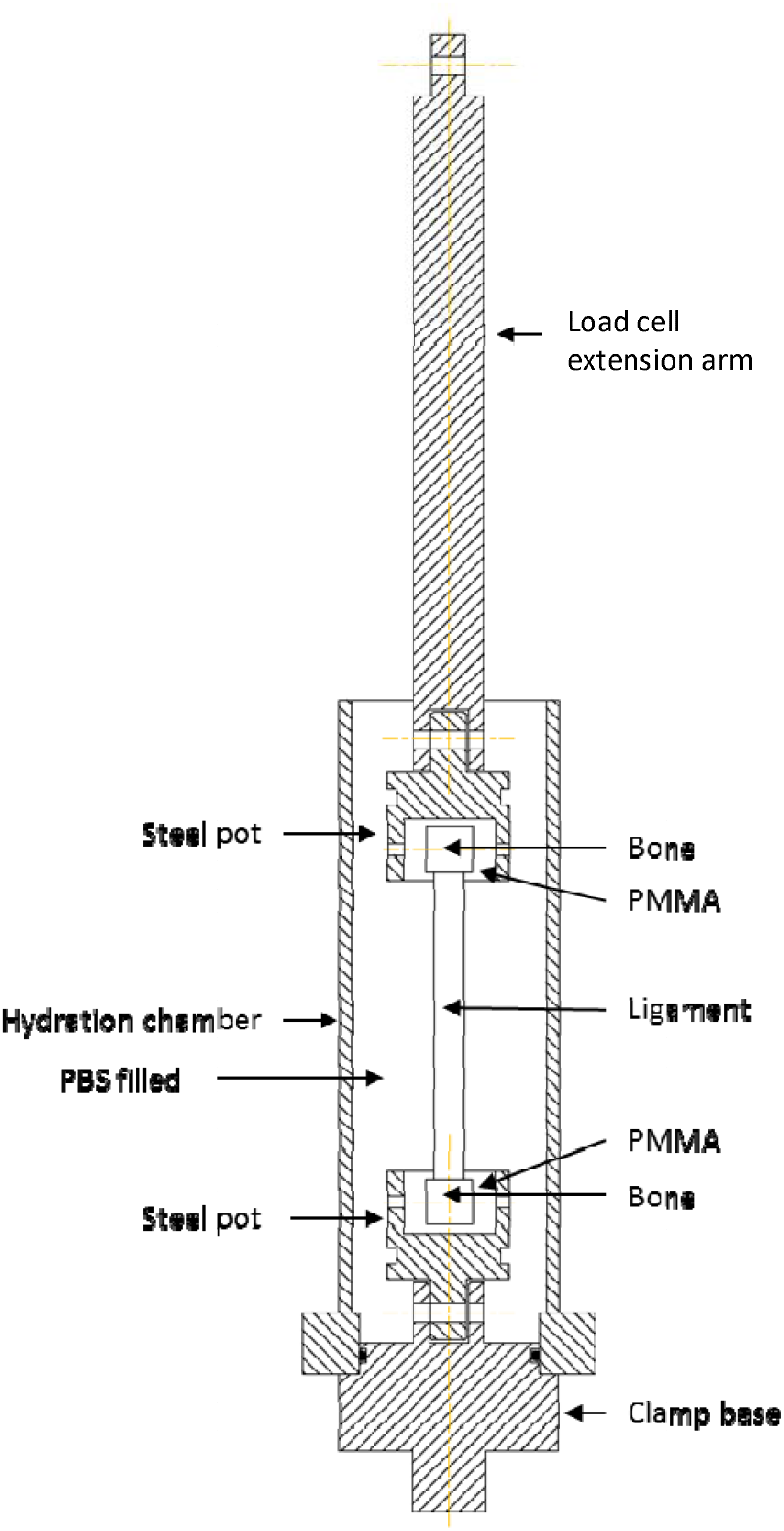
Schematic illustration of the custom-made rig for tensile testing of human knee joint ligaments. The bone ends of the ligament were secured by potting them into a polymethylmethacrylate (PMMA) holder. Ligaments were encased into a watertight chamber filled with phosphate buffer saline (PBS) to maintain tissue hydration during mechanical tests.

### Tensile Testing Protocol

Using a uniaxial tensile testing machine (Instron 3366, Buckinghamshire, UK) with a 5000 N load cell (Instron 2519), a 1 N preload was applied and all ligaments underwent ten preconditioning cycles at 10 mm/min with a load of 1 to 40 N, which provides a stable and repeatable viscoelastic response (Momersteeg et al., 1995). Loading was then set to zero and ligaments were loaded to failure at 500 mm/min. A fast strain rate was chosen over slow stain rates to mimic physiological loading (Noyes & Grood, 1976; Sharma et al., 2008; Bersini, Sansone & Frigo, 2016) and replicate a realistic injury environment (Robinson, Bull & Amis, 2005). In addition, faster strain rates improve the chances of the ligament rupturing mid-substance as opposed to a bony avulsion (Noyes & Grood, 1976).

### Material Properties

The bone-ligament-bone specimens were mechanically tested and analyzed to collate multiple material property data (Fig. 3). Linear parameters were calculated from the last data point on the linear slope of the curve, whereas failure parameters were calculated from the attained maximum load, the parameters were force, elongation, stress, strain, secant modulus, tangent modulus (the slope between maximum and sub-maximum points of the linear region of the load-extension curve) and stiffness (Fig. 3).

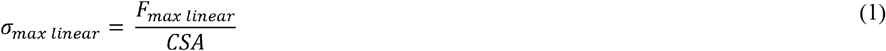

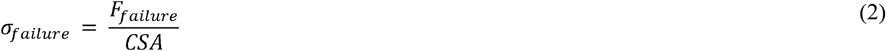

where *σ_max linear_* and (*σ_failure_* are stresses (MPa) at the maximum linear and failure points, *F_max linear_* and *F_failure_* are forces (N) at the maximum linear and failure points, and *CSA* is cross-sectional area (mm^2^).

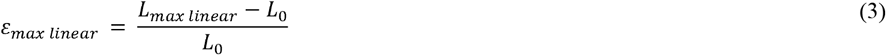

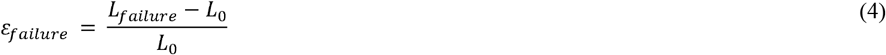

where *ε_max linear_* and *ε_failure_* are strains (%) at the maximum linear and failure points, *L_max linear_* and *L_failure_* are lengths (mm) at the maximum linear and failure points, and *L*_0_ is original length (mm) of the ligament.

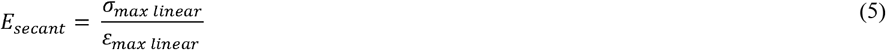

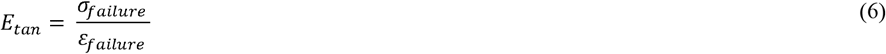

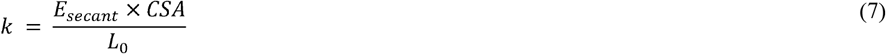

where *E_secant_* is secant modulus (MPa), *E_tan_* is tangent modulus (MPa) at the maximum linear region of the load-extension curve, and *k* is ligament stiffness (N/mm).

**Fig. 3.**
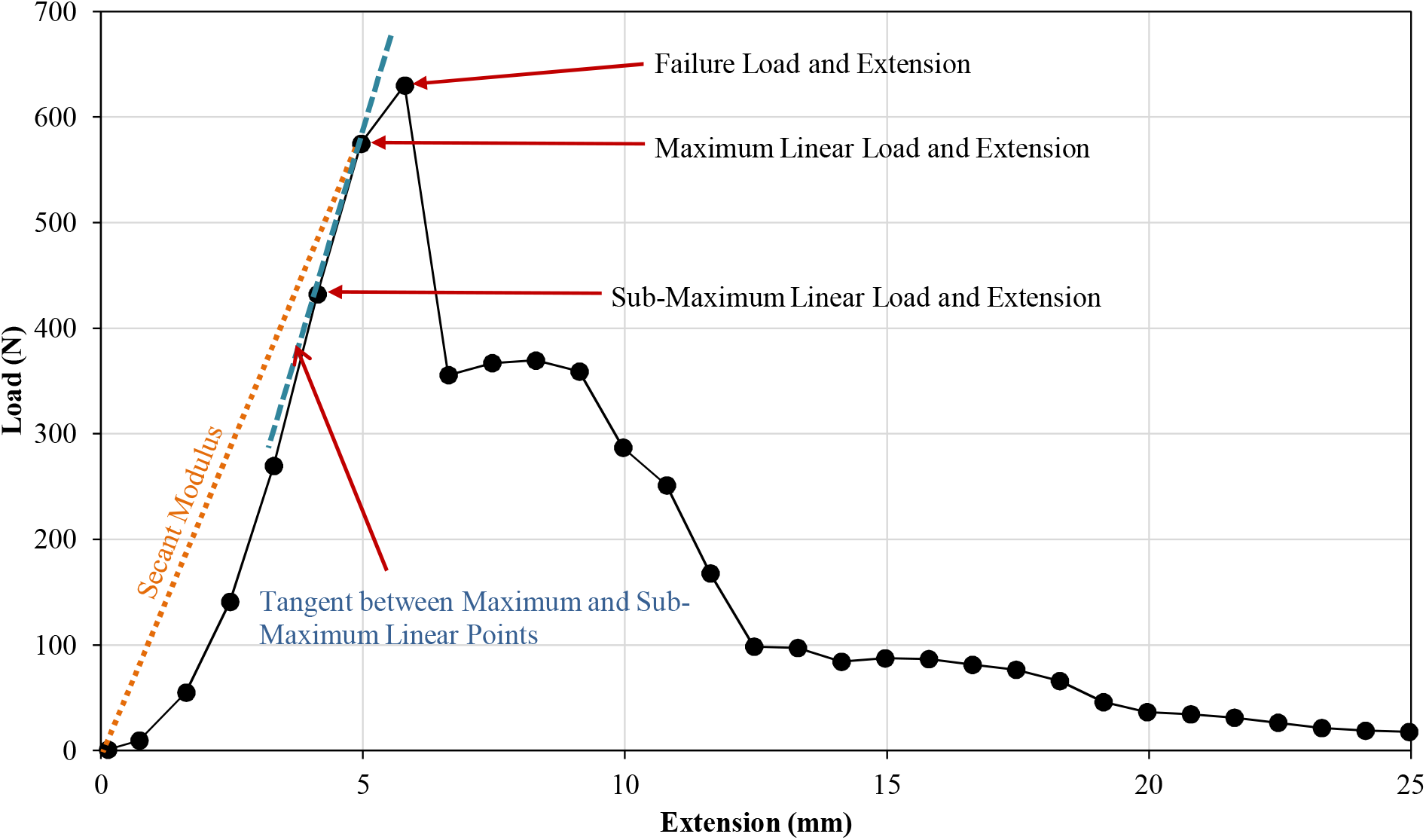
Example of a load-extension curve, showing failure load and elongation point, maximum and sub-maximum linear loads in Newtons (N), and extensions in millimetres (mm) human knee joint ligaments. The figure also highlights the tangent modulus between maximum and sub-maximum linear points and the secant modulus.

### Statistical Analysis

Kruskal-Wallis one-way ANOVA was conducted to compare mean differences of ligament material properties between young healthy (≤60 years old, ICRS grade 0), young OA (≤60 years old, ICRS grade 1-4) and old OA (>60, ICRS grade 1-4) cohorts. Ligament tensile properties were correlated with increasing age and grade of OA using a Kendall’s Tau-b (*τ*_bc_) correlation coefficient. The material properties included in the analyses were: linear force, linear stress, linear strain, failure load, failure stress, failure strain, secant modulus, tangent modulus, and stiffness. Both of the statistical analyses were performed in SPSS (SPSS software, Version 22.0, SPSS, Inc., Chicago, IL).

## Results

### Specimens

ACL (n = 12), PCL (n = 12), MCL (n = 12), and LCL (n = 12) specimens were obtained from twelve human cadavers. One MCL specimen from a young (37 years old) healthy donor was visually determined as severely abnormal and data from two MCL specimens from ICRS grade 1 donors were unable to be retained (Supplementary Materials (Fig. S1)). Hence, they were excluded from statistical analyses.

The ICRS grading for all 12 cadaveric knees were given and reported in Table 1 which is the same as those reported in our previous work (Peters et al., 2018b). Three knees were given ICRS grade 0 (age: 31, 37 and 43 years old), another three knees with ICRS grade 1 (age: 49, 51 and 86 years old), two knees with ICRS grade 2 (age: 58 and 79 years old), three knees with ICRS grade 3 (age: two 72 and 88 years old) and one knee with ICRS grade 4 (age: 80 years old).

**Table 1:** Anterior (ACL) and posterior (PCL) cruciate ligaments, medial (MCL) and lateral (LCL) collateral ligaments material property data for all cadavers. ABREVIATIONS: OA ICRS, osteoarthritis International Cartilage Repair Society; *L*_0_, original length of the ligament; *CSA,* cross-sectional area; *F,* force or load; *σ*, stress; *ε*, strain at the maximum point of the linear region (_max linear_) and failure (_failure_) point of the load-extension curve; ***E_secant_***, secant modulus; ***E_tan_***, tangent modulus; *k,* stiffness; I, insertion; MS, mid-substance; BA, bony avulsion.

| Age (years) | Sex | OA ICRS Grade | Ligament | $L_0$ (mm) | CSA (mm<sup>2</sup>) | $F_{\max \text{ linear}}$ (N) | $\sigma_{\max \text{ linear}}$ (MPa) | Max Linear Extension (mm) | $\epsilon_{\max \text{ linear}}$ (%) | $E_{\text{secant}}$ (MPa) | $E_{\text{tan}}$ (MPa) | $F_{\text{failure}}$ (N) | $\sigma_{\text{failure}}$ (MPa) | Failure extension (mm) | $\epsilon_{\text{failure}}$ (%) | $k$ (N/mm) | Failure Site |
| --- | --- | --- | --- | --- | --- | --- | --- | --- | --- | --- | --- | --- | --- | --- | --- | --- | --- |
| 31 | Female | 0 | ACL | 40 | 63.80 | 574.78 | 9.01 | 4.96 | 12.40 | 29.06 | 171.04 | 629.94 | 9.87 | 5.79 | 14.48 | 46.36 | I |
|  |  |  | PCL | 36 | 86.53 | 624.66 | 7.22 | 4.34 | 12.05 | 21.57 | 238.72 | 930.27 | 10.75 | 8.50 | 23.62 | 51.84 | MS |
|  |  |  | LCL | 62 | 17.10 | 477.63 | 27.93 | 10.98 | 17.71 | 97.79 | 95.80 | 545.90 | 31.92 | 11.81 | 19.05 | 26.98 | MS |
|  |  |  | MCL | 103 | 24.51 | 332.35 | 13.56 | 6.74 | 6.54 | 213.56 | 85.96 | 438.48 | 17.89 | 8.40 | 8.16 | 50.82 | MS |
| 37 | Female* | 0 | ACL | 30 | 48.46 | 1345.42 | 27.76 | 6.75 | 22.50 | 37.03 | 239.08 | 1554.16 | 32.07 | 8.42 | 28.05 | 59.81 | BA |
|  |  |  | PCL | 30 | 48.82 | 1107.59 | 22.69 | 9.72 | 32.41 | 21.00 | 293.42 | 1336.49 | 27.37 | 10.56 | 35.19 | 34.17 | BA |
|  |  |  | LCL | 55 | 71.75 | 280.88 | 3.91 | 7.89 | 14.34 | 15.02 | 70.23 | 479.73 | 6.69 | 11.22 | 20.40 | 19.59 | I |
|  |  |  | MCL | 40 | 18.19 | 33.21 | 1.83 | 4.06 | 10.15 | 7.19 | 14.32 | 47.93 | 2.63 | 4.90 | 12.24 | 3.27 | MS |
| 43 | Female | 0 | ACL | 32 | 71.27 | 467.52 | 6.56 | 5.78 | 18.08 | 11.61 | 126.42 | 576.94 | 8.09 | 9.95 | 31.10 | 25.86 | MS |
|  |  |  | PCL | 30 | 76.53 | 369.20 | 4.82 | 6.83 | 22.78 | 6.35 | 73.99 | 1220.92 | 15.95 | 15.17 | 50.56 | 16.21 | MS |
|  |  |  | LCL | 61 | 12.83 | 81.49 | 6.35 | 4.39 | 7.19 | 53.89 | 34.31 | 426.52 | 33.25 | 11.05 | 18.12 | 11.33 | MS |
|  |  |  | MCL | 108 | 28.29 | 343.16 | 12.13 | 11.16 | 10.34 | 126.76 | 75.41 | 727.51 | 25.72 | 19.50 | 18.05 | 33.20 | MS |
| 49 | Male | 1 | ACL | 40 | 34.68 | 229.22 | 6.61 | 6.87 | 17.17 | 15.40 | 120.79 | 340.82 | 9.83 | 9.37 | 23.42 | 13.35 | MS |
|  |  |  | PCL | 44 | 84.79 | 777.84 | 9.17 | 5.96 | 13.55 | 29.79 | 229.97 | 963.21 | 11.36 | 9.29 | 21.12 | 57.41 | MS |
|  |  |  | LCL | 52 | 39.91 | 279.40 | 7.00 | 15.88 | 30.55 | 11.92 | 67.32 | 438.52 | 10.99 | 21.72 | 41.76 | 9.15 | MS |
|  |  |  | MCL | 101 | 22.37 | 137.53 | 6.15 | 10.92 | 10.81 | 57.44 | 35.88 | 552.09 | 24.68 | 17.59 | 17.41 | 12.72 | MS |
| 51 | Male | 1 | ACL | 28 | 53.64 | 249.76 | 4.66 | 4.89 | 17.48 | 7.46 | 53.79 | 504.97 | 9.41 | 12.39 | 44.27 | 14.29 | MS |
|  |  |  | PCL | 34 | 59.24 | 378.24 | 6.38 | 4.89 | 14.38 | 15.10 | 120.23 | 1052.61 | 17.77 | 15.72 | 46.24 | 26.31 | I |
|  |  |  | LCL | 47 | 45.15 | 341.77 | 7.57 | 7.04 | 14.97 | 23.77 | 81.27 | 419.25 | 9.29 | 15.37 | 32.70 | 22.83 | I |
|  |  |  | MCL | 114 | 41.18 | 279.31 | 6.78 | 7.32 | 6.42 | 120.38 | 61.30 | 354.33 | 8.61 | 13.99 | 12.27 | 43.48 | MS |
| 58 | Male | 2 | ACL | 41 | 95.79 | 169.90 | 1.77 | 6.97 | 17.00 | 4.28 | 55.47 | 664.31 | 6.93 | 14.47 | 35.29 | 10.00 | MS |
|  |  |  | PCL | 46 | 98.67 | 980.12 | 9.93 | 6.89 | 14.99 | 30.49 | 220.75 | 1380.78 | 13.99 | 10.23 | 22.23 | 65.41 | I |
|  |  |  | LCL | 58 | 36.03 | 388.10 | 10.77 | 5.88 | 10.14 | 61.60 | 103.30 | 629.39 | 17.47 | 9.22 | 15.89 | 38.27 | MS |
|  |  |  | MCL | 127 | 28.80 | 150.06 | 5.21 | 7.85 | 6.18 | 107.01 | 30.76 | 514.05 | 17.85 | 14.52 | 11.43 | 24.27 | MS |
| 72 (1) | Male | 3 | ACL | 34 | 49.75 | 569.38 | 11.44 | 3.96 | 11.66 | 33.39 | 197.08 | 808.34 | 16.25 | 5.63 | 16.56 | 48.85 | MS |
|  |  |  | PCL | 41 | 62.51 | 344.93 | 5.52 | 7.67 | 18.71 | 12.09 | 79.10 | 524.83 | 8.40 | 11.84 | 28.88 | 18.43 | BA |
|  |  |  | LCL | 60 | 66.07 | 251.52 | 3.81 | 9.87 | 16.46 | 13.88 | 51.08 | 379.68 | 5.75 | 15.71 | 26.18 | 15.28 | MS |
|  |  |  | MCL | 121 | 33.41 | 235.88 | 7.06 | 8.33 | 6.88 | 124.16 | 71.24 | 458.47 | 13.72 | 12.49 | 10.32 | 34.28 | I |
| 72 (2) | Male | 3 | ACL | 29 | 101.84 | 265.33 | 2.61 | 2.35 | 8.10 | 9.33 | 189.12 | 613.77 | 6.03 | 4.85 | 16.72 | 32.76 | I |
|  |  |  | PCL | 31 | 91.34 | 1059.28 | 11.60 | 5.03 | 16.23 | 22.15 | 321.15 | 1432.70 | 15.69 | 6.70 | 21.60 | 65.28 | MS |
|  |  |  | LCL | 68 | 44.46 | 282.10 | 6.35 | 7.18 | 10.55 | 40.89 | 66.36 | 352.48 | 7.93 | 11.34 | 16.68 | 26.73 | MS |
|  |  |  | MCL | 110 | 58.68 | 120.41 | 2.05 | 3.64 | 3.31 | 68.15 | 62.58 | 227.10 | 3.87 | 13.64 | 12.40 | 36.35 | MS |
| 79 | Male | 2 | ACL | 32 | 37.78 | 145.15 | 3.84 | 7.43 | 23.21 | 5.30 | 46.39 | 187.78 | 4.97 | 10.76 | 33.63 | 6.25 | I |
|  |  |  | PCL | 32 | 70.34 | 356.25 | 5.06 | 5.61 | 17.54 | 9.24 | 100.22 | 642.60 | 9.14 | 12.28 | 38.37 | 20.31 | MS |
|  |  |  | LCL | 62 | 18.98 | 532.16 | 28.03 | 8.81 | 14.21 | 122.29 | 87.79 | 628.01 | 33.08 | 12.15 | 19.59 | 37.44 | I |
|  |  |  | MCL | 120 | 39.57 | 264.96 | 6.70 | 17.86 | 14.89 | 53.98 | 51.74 | 326.72 | 8.26 | 19.53 | 16.27 | 17.80 | I |
| 80 | Male | 4 | ACL | 38 | 74.89 | 139.35 | 1.86 | 5.67 | 14.91 | 4.74 | 77.25 | 373.94 | 4.99 | 9.00 | 23.68 | 9.35 | MS |
|  |  |  | PCL | 35 | 154.25 | 65.96 | 0.43 | 4.89 | 13.98 | 1.07 | 29.79 | 261.57 | 1.70 | 15.73 | 44.94 | 4.72 | I |
|  |  |  | LCL | 74 | 50.01 | 250.44 | 5.01 | 11.65 | 15.74 | 23.54 | 59.90 | 445.35 | 8.91 | 21.65 | 29.26 | 15.91 | BA |
|  |  |  | MCL | 116 | 27.62 | 399.01 | 14.45 | 7.68 | 6.62 | 253.25 | 73.99 | 492.60 | 17.84 | 12.68 | 10.93 | 60.29 | MS |
| 86 | Female** | 1 | ACL | 30 | 24.98 | 84.17 | 3.37 | 2.57 | 8.58 | 11.78 | 58.95 | 134.33 | 5.38 | 5.91 | 19.69 | 9.81 | I |
|  |  |  | PCL | 43 | 66.54 | 67.81 | 1.02 | 4.68 | 10.89 | 4.02 | 40.68 | 281.04 | 4.22 | 13.85 | 32.21 | 6.23 | I |
|  |  |  | LCL | 60 | 14.17 | 89.69 | 6.33 | 6.91 | 11.52 | 32.98 | 33.56 | 263.26 | 18.58 | 13.58 | 22.63 | 7.79 | MS |
| 88 | Male | 3 | ACL | 33 | 64.32 | 144.40 | 2.25 | 5.18 | 15.68 | 4.72 | 58.76 | 277.47 | 4.31 | 9.34 | 28.31 | 9.21 | I |
|  |  |  | PCL | 34 | 95.25 | 212.68 | 2.23 | 6.33 | 18.60 | 4.08 | 91.30 | 414.07 | 4.35 | 9.66 | 28.41 | 11.43 | I |
|  |  |  | LCL | 58 | 24.78 | 101.97 | 4.12 | 6.44 | 11.10 | 21.50 | 31.79 | 337.54 | 13.62 | 13.94 | 24.03 | 9.19 | I |
|  |  |  | MCL | 120 | 35.55 | 232.72 | 6.55 | 9.55 | 7.96 | 98.69 | 41.48 | 444.18 | 12.49 | 17.05 | 14.21 | 29.24 | MS |

\* Donor had a severely abnormal MCL and was not included in the statistical analysis.
\*\* The MCL from this doner could not be retained for mechanical tests.

### Cross-Sectional Area & Length Measurements

The cross-sectional areas of the ACLs, PCLs, LCLs and MCLs were in the range of 24.98 to 101.84, 48.82 to 154.25, 12.83 to 71.75, and 18.19 to 58.68 mm^2^, respectively. The length of the ACLs, PCLs, LCLs and MCLs were in the range of 28 to 41, 30 to 46, 47 to 74, and 40 to 127 mm, respectively. The cross-sectional areas and lengths of individual ligaments for each donor are reported in Table 1.

### Correlation with Age

Increasing age resulted in statistically significant negative correlations with ACL linear force (*τ_b_ = −0.63, p = 0.01*), linear stress (*τ_b_ = −0.47, p = 0.03*), linear extension (*τ_b_ = −0.44, p = 0.05*), failure force (*τ_b_ = −0.50, p = 0.02*) and failure stress (*τ_b_ = −0.53, p = 0.02*) (Fig. 4). No statistically significant correlation was found between age and ACL linear strain, secant and tangent modulus, failure strain and stiffness (Supplementary Materials (Table S4 and ACL_AllStat.xls)).

**Fig. 4.**
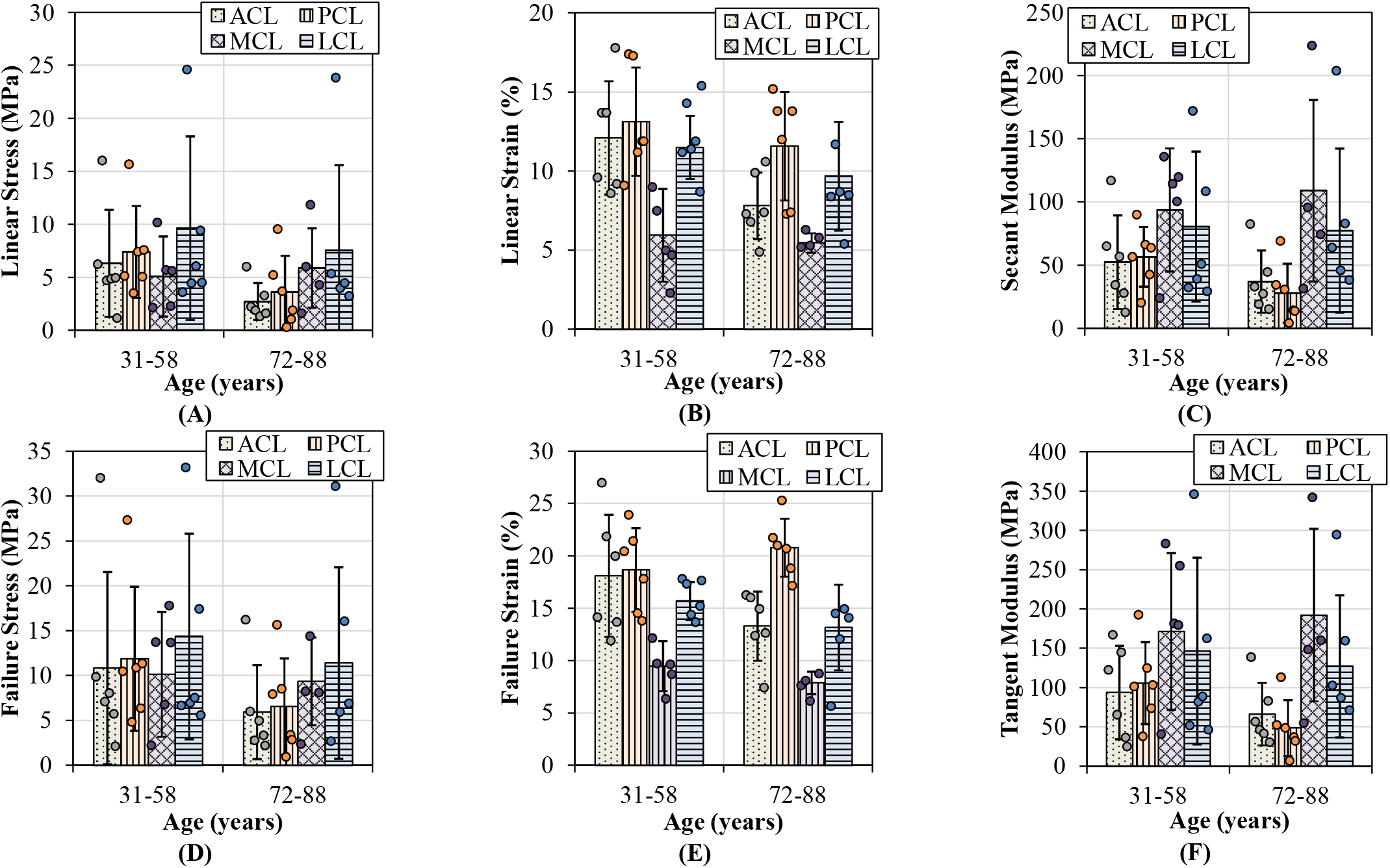
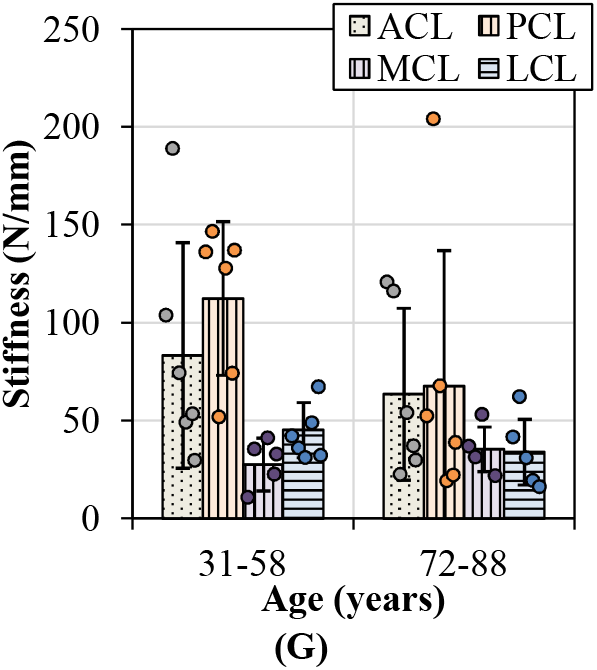
Tensile parameters for the anterior (ACL) and posterior (PCL) cruciate ligaments, and lateral (LCL) and medial (MCL) collateral ligaments across two age groups (31-58 and 72-88 years old). Error bars represent standard deviation. (A) Linear stress and (B) linear strain were utilized to determine (C) secant modulus. (D) and (E) demonstrates the maximum stresses and strains that resulted in ligament failures. (F) This figure shows tangent modulus of the ligaments between the two age groups at the maximum linear region of load-extension curves. (G) This figure documents the change in stiffness of the ligaments with age.

Increasing age showed statistically significant negative correlations with PCL secant modulus (*τ_b_ = −0.50, p = 0.02*) and tangent modulus (*τ_b_ = −0.53, p = 0.02).* No statistically significant correlations were found between age and the rest of the PCL tensile parameters (Supplementary Materials (Table S4 and PCL_AllStat.xls)).

There were no statistically significant correlations between age and MCL tensile parameters (Supplementary Materials (MCL_AllStat.xls)). Only LCL stiffness showed statistically significant negative correlation with age (*τ_b_ = −0.44, p = 0.05*) and no additional significant correlations were found for the LCL tensile properties (Supplementary Materials (Table S4 and LCL_AllStat.xls)).

Detailed correlation of age with material properties of the four ligaments are reported in the Supplementary Materials (Table S3, Table S4 and Fig. S3).

### Correlation with Osteoarthritis

Increasing OA grade showed statistically significant negative correlation with ACL linear force (*τ_b_ = −0.46, p = 0.05*), linear stress (*τ_b_ = −0.53, p = 0.02*), linear extension (*τ_b_ = −0.59, p = 0.01*) and linear strain (*τ_b_ = −0.5, p = 0.03).* However, the correlations between OA and the rest of the ACL tensile parameters were not statistically significant (Supplementary Materials (Table S4 and ACL_AllStat.xls)).

There were no statistically significant correlations between OA grade and PCL and MCL tensile parameters. Only LCL failure stress showed statistically significant negative correlation between OA grade and LCL failure stress (*τ_b_ = −0.46, p = 0.05*) and the rest of the LCL tensile parameters were not statistically significant (Supplementary Materials (PCL_AllStat.xls, MCL_AllStat.xls, and LCL_AllStat.xls)).

Detailed correlation of OA with material properties of the four ligaments are reported in the Supplementary Materials (Table S3, Table S4 and Fig. S3).

## Discussion

This is the first *ex vivo* study to quantify of the effects of aging and OA on the material properties of the four primary knee ligaments from the same cadaveric joints within a wide span of age (31 to 88 years old) and OA grade (ICRS 0 to 4). Our results showed statistically significant negative correlations with ACL linear and failure forces, stresses and extensions, PCL secant and tangent modulus and LCL stiffness (Supplementary Materials (Table S4)). Similarly, increasing OA grade resulted in statistically significant negative correlation with ACL linear forces, stresses, extension, and strains and LCL failure stress (Supplementary Materials (Table S4)). Change in age or OA grade did not have a significant correlation with the MCL material parameters (Supplementary Materials (Table S4)). This data is vital for understanding joint mechanics and it can provide an insight into the progression of OA as a whole-joint disease as well as the effects of aging, notably because bone and cartilage mechanical properties for these specific human cadavers have already been reported in our previous study (Peters et al., 2018b).

Failure loads previously reported across any age category span two orders of magnitude between 495 to 2160 N in the ACL, 258 to 1620 N in the PCL, 194 to 534 N in the MCL and 376 N in the LCL (Noyes & Grood, 1976; Trent, Walker & Wolf, 1976; Woo et al., 1991; Race & Amis, 1994; Harner et al., 1995; Chandrashekar et al., 2006). Furthermore, previous studies also reported stiffness values which ranged between 124 to 308 N/mm in the ACL, 57 to 347 N/mm in the PCL, 70 N/mm in the MCL and 59 N/mm in the LCL, where values reported for failure load and stiffness in the current study fall within the previously reported range (Noyes & Grood, 1976; Trent, Walker & Wolf, 1976; Woo et al., 1991; Race & Amis, 1994; Harner et al., 1995; Chandrashekar et al., 2006). Previous research has indicated a decrease in the ACL failure load with increasing age, which is consistent with the current study (Fig. 4D). Age based differences show ACL failure loads of up to 2160 N amongst younger donors (22 to 35 years), 1503 N in middle-aged donors (40 to 50 years) and 658 N amongst older donors (60 to 97 years), although degeneration of joint integrity was not indicated (Woo et al., 1991).

The current research showed a decrease in the failure strain of all four knee ligaments with the development of OA (Fig. 5E and Supplementary Materials (Fig. S3)). The ACL in healthy knees showed higher linear and failure stresses and strains, secant and tangent modulus and stiffness when compared to those with OA (Fig. 5). The influence of OA has previously been investigated in animal models and a reduction in tensile properties of the rat ACL was reported, ten weeks after collagen-induced arthritis. Ultimate failure load reduced by 25.1% and stiffness by 38.0% when compared to controls (Nawata et al., 2001). Despite a lack of knee joint material properties in the literature associated to OA in humans, previous research has found that between 39 to 78% of patients with OA have a degenerated ACL (Allain, Goutallier & Voisin, 2001; Cushner et al., 2003; Lee et al., 2005; Mullaji et al., 2008; Watanabe et al., 2011), and between 7 to 80% have a degenerated PCL (Nelissen & Hogendoorn, 2001; Stubbs et al., 2005; Mullaji et al., 2008). Such degeneration is consistent with the decrease in the tensile properties of the four knee joint ligaments observed in our current study.

**Fig. 5.**
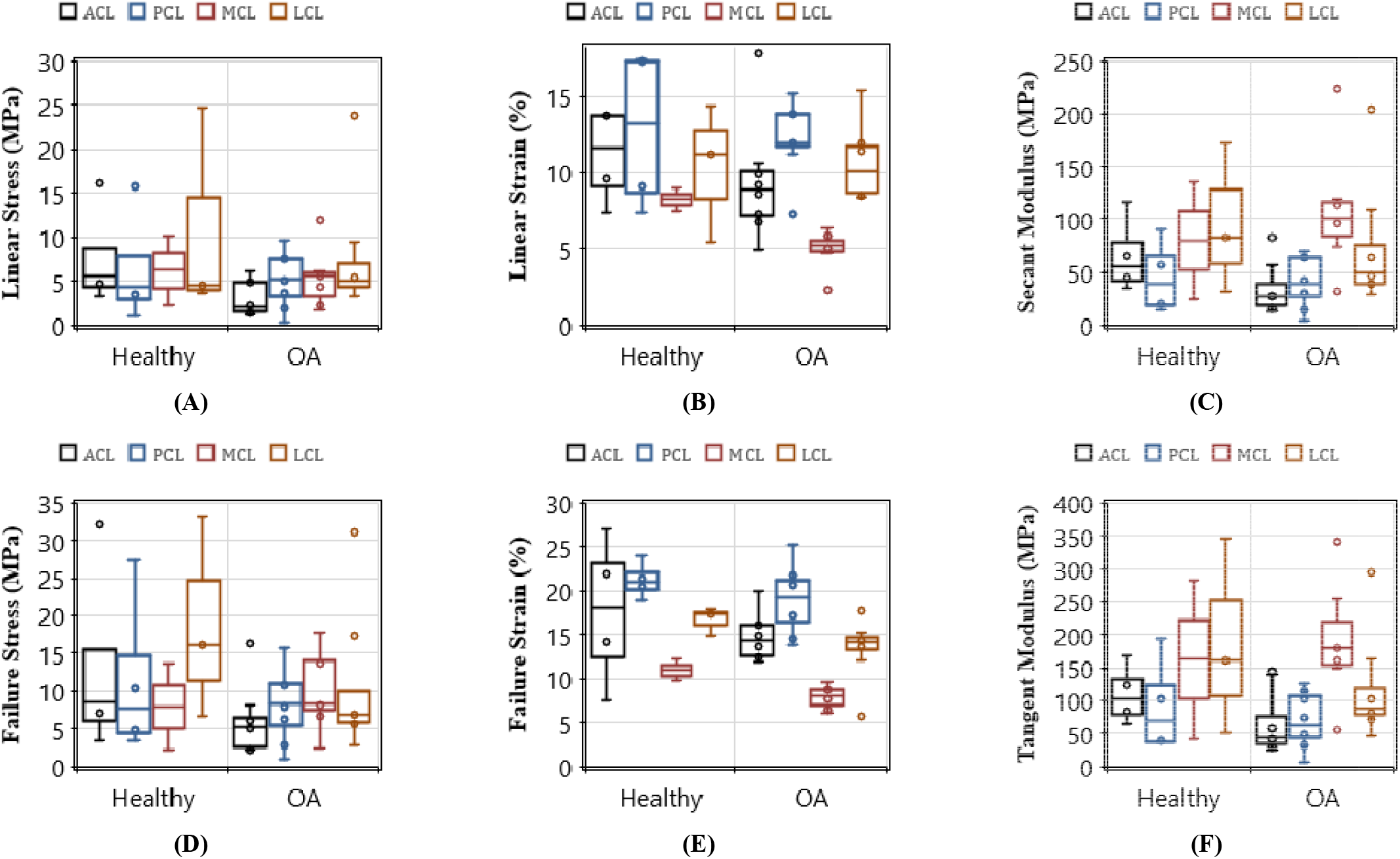
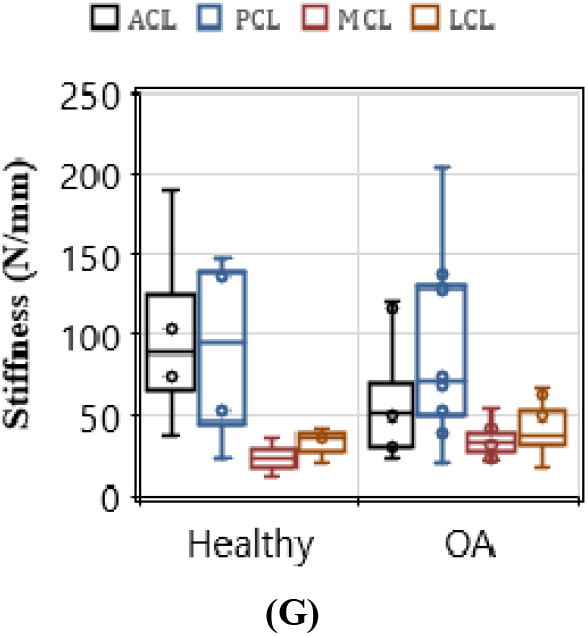
Comparisons of tensile properties of the anterior (ACL) and posterior (PCL) cruciate ligaments, and lateral (LCL) and medial (MCL) collateral ligaments between healthy and osteoarthritic (OA) groups. Healthy groups were defined by International Cartilage Repair Society (ICRS) grade 0 and osteoarthritic (OA) was defined by ICRS grade 1-4. (A) Linear stress and (B) linear strain were utilized to determine (C) secant modulus. (D) and (E) demonstrates the maximum stresses and strains that resulted in ligament failures. (F) This figure shows tangent modulus of the ligaments between the healthy and OA groups at the maximum linear region of load-extension curves. (G) This figure documents ligament stiffness values across ligaments and between the healthy and OA groups.

Currently, it is challenging to separate the effects of OA and aging as they often happen concurrently. With only 12 cadavers and five groups of ICRS grades (0 to 4), it was challenging to statistically attribute changes in ligament tensile properties to both age and OA as related parameters, particularly when also accounting for sex (see further discussion below). This is a limiting factor of the current study; however, to understand the effect of age and OA as individual parameters, trends were analyzed from the data presented in Table 1. The trends suggest that in younger donors the ACL and PCL material properties were reduced in the OA knees compared to those in the healthy knees (Supplementary Materials (Fig. S4)). This suggests that even mild OA in younger donors affects the material properties which is further exacerbated with advancing age and OA.

It is currently unknown whether ligament injury initiates the early-onset of OA, or whether OA is indeed a whole-joint disease impairing the integrity of associated tissues including ligaments (Poole, 2012). In our previous study on the same human cadaveric knees, we found statistically significant correlations between changes in material properties of cartilage and subchondral bone with age and OA grade (Peters et al., 2018b). Similarly, the data in the current study for the same cadavers showed alterations of ligament tensile properties because of OA. It is possible that ligament degeneration or injury occurs in the first instance and this can subsequently lead to the initiation and progression of knee OA (Gianotti et al., 2009). Since the primary function of knee ligaments is to provide stability to the knee joint (Harner et al., 1995; Woo et al., 2006), any changes to the ligaments’ structure can alter the load distribution in the knee joint (Moore & Burris, 2015). The knee cadavers in this study showed that OA degeneration affected the medial more than the lateral compartments of the bones (Peters et al., 2018b). The difference in degeneration between the lateral and medial compartments of the knee joint could be a result of unbalanced load distribution that was caused by changes in the ligament material properties because of OA. Our data also showed that PCL tensile properties most commonly decreased between younger donors with the presence of OA and older donors with the presence of OA (Supplementary Materials (Fig. S4)). This suggests that changes to this ligament are more evident with advanced aging. The reduction in the measured tensile parameters of the ACL during aging and disease progression may be attributed to the relatively high forces experienced during walking. Studies show a consensus that peak force experienced by the ACL occurs at contralateral toe off during the stance phase of the gait cycle, up to 3.5 times body weight (Morrison, 1970; Collins & O’Connor, 1991; Shelburne et al., 2004). In particular, these high ACL kinematic forces may be consistent with the widely reported histological degeneration of the ACL in the presence of disease (Mullaji et al., 2008), suggesting high habitual forces could influence subsequent degeneration observed. Peak force of the PCL has also been reported up to 0.2 to 0.6 times body weight during walking (Morrison, 1970; Collins & O’Connor, 1991). There is evidence showing that appropriate exercise training strengthens ligaments and knee joint mechanics (Tipton et al., 1975; Salem et al., 2000; Ng, 2002; Ferri et al., 2003). However, people exercise less as they age hence increasing their risk of ligament degeneration (Daley & Spinks, 2000). Decreased capacity of the knee ligaments to resist motion due to reduced mechanical strength may alter contact forces of the joint, potentially causing increased loading on the medial femoral condyle and contributing to the preferential medial development of OA (Pelletier et al., 2007; Lohmander et al., 2007).

Further limitations of the current study, aside from a low sample number, include varying donor demographics, such as sex, which is known to affect tensile properties and likelihood of knee ligament injury. It was found that ACLs in young human females had 22.49% lower Young’s modulus, and 8.3% and 14.3% lower failure strain and stress respectively, when compared to ACLs in young human males (Chandrashekar et al., 2006). These differences can be partially attributed to the physically smaller size of the female ACL, which can in turn be linked to higher rates of ACL injuries in female athletes (Anderson et al., 2001; Chandrashekar, Slauterbeck & Hashemi, 2005). Females are also known to be at a greater risk of knee OA than males (Hame & Alexander, 2013). Again, due to low sample numbers, this study was unable to separate ligaments by sex for statistical analyses.

Finally, the current study may be limited by testing ligaments as whole bone-ligament-bone specimens along their loading axis. It has previously been acknowledged that ligaments may be best divided into their fiber bundles in order to recruit fibers to their maximal potential and eliminate any slack due to orientation (Woo et al., 1991; Race & Amis, 1994). Significant differences have been reported between the anterior and posterior fibers of the ACL (Butler et al., 1992) and PCL (Race & Amis, 1994; Harner et al., 1995) suggesting that fibers play different roles in the stabilization of the knee joint (Race & Amis, 1994); although ligaments naturally work as one functional unit. Such global approaches have previously been used in the representation of ligaments in finite element models as one functional unit (Readioff et al., 2020a). However, due to the lack of data on all four ligaments from the same donor (and in certain cases the same demographic or disease conditions of donor) in the literature, material properties have often been applied globally in finite element models, where values for one ligament are replicated for all others (Blankevoort & Huiskes, 1991; Li, Lopez & Rubash, 2001; Wang, Fan & Zhang, 2014; Kazemi & Li, 2014). In some instances, tendon material properties have been used (Kazemi et al., 2011; Wang, Fan & Zhang, 2014; Kazemi & Li, 2014). Sensitivity analysis showed that varying intrinsic ligament material properties alters the internal and external rotation of the tibia-femoral joint, patella tilt and peak contact stress (Dhaher, Kwon & Barry, 2010). As such, the data in this study combined with cartilage and bone data in our previous study (Peters et al., 2018b) allows future research to apply a subject- or cohort-specific approach to computational modelling of the human knee joint to improve accuracy and predictive behavior patterns of ligaments.

Furthermore, the knowledge of baseline material properties of all four ligaments from healthy donors can be used to replicate ligaments by developing more biologically representative synthetic materials for the repair and replacement following injury or degeneration (Yang, Rothrauff & Tuan, 2013; Ratcliffe et al., 2015). The data collected in this study, therefore, provides insight into not only the healthy range for these parameters but also how they change concurrently with surrounding ligaments during aging and disease.

## Conclusions

This research is the first to correlate alterations in tensile properties of the four major human knee ligaments to aging and OA. We confirmed the findings of previous research that the ACL tensile properties decrease with age and OA. The results also provided new evidence that the PCL tangent and secant modulus decrease with increasing age. Although not statistically proven, the MCL and LCL showed some changes with increasing age and OA grade, which has not previously been demonstrated. These data, along with our previously reported data on bone and cartilage material properties for the same cadavers supports current research stating that OA is a whole-joint disease impairing many peri-articular tissues within the knee. The material properties for the four major knee ligaments in the twelve cadavers can be combined with their corresponding subchondral and trabecular bones and articular cartilage for future subject-specific applications including the development of computational models and OA diagnostics.

## Acknowledgements

This project was funded by BBSRC (Research Grant: BB/J014516/1) and the School of Engineering, University of Liverpool.

The authors would like to thank Mr Phil Jackson and the staff at the Newcastle Surgical Training Centre, who supported obtainment of cadaveric specimens, and Mr Lee Moore and the staff at the Veterinary Training Suite, Institute of Veterinary Science, University of Liverpool, for the use of surgical tools. We also thank Prof. Ahmed Elsheikh for providing access to the experimental facilities at the School of Engineering, University of Liverpool.

