## Supplemental Materials for "Biomechanics of aging and osteoarthritic human knee ligaments"

**Supplementary Materials**

**Table S1**: International Cartilage Repair Society (ICRS) Grading.

| ICRS Grade | Description |
| --- | --- |
| 0 (Normal) | No lesions fissures or cracks. |
| 1 (Nearly Normal) | Superficial lesions. Soft indentation and/or superficial fissures and cracks. |
| 2 (Abnormal) | Lesions extending down to <50% of cartilage depth. |
| 3 (Severely Abnormal) | Cartilage defects extending down >50% of cartilage depth as well as down to calcified layer and down to but not through the subchondral bone. Blisters are included in this Grade. |
| 4 (Severely Abnormal) | Cartilage defects extending down >75% of cartilage depth as well as down to calcified layer and through the subchondral bone. Blisters are included in this Grade. |

**Table S2**: Cadaver demographics.

|  | Age | Gender | Race | Height (cm) | Weight (kg) | BMI | Cause of Death |
| --- | --- | --- | --- | --- | --- | --- | --- |
| Cadaver 1 | 31 | Female | Not known | 172.7 | 47.2 | 15.81 | Cardiac arrest |
| Cadaver 2 | 37 | Female | White | 160.0 | 79.4 | 31.00 | Intracerebral haemorrhage; Severe hypertension |
| Cadaver 3 | 43 | Female | White | 170.2 | 64.4 | 22.24 | Metastatic cervical carcinoma |
| Cadaver 4 | 49 | Male | White | 175.3 | 58.5 | 19.05 | Not known |
| Cadaver 5 | 51 | Male | White | 182.9 | 104.3 | 31.19 | Cardiac arrhythmia; Coronary artery disease |
| Cadaver 6 | 58 | Male | White | 188.0 | 84.8 | 24.01 | Gunshot wound of head and neck |
| Cadaver 7 | 72 | Male | Puerto Rican | 162.6 | 70.3 | 26.60 | Atherosclerotic heart disease of native coronary |
| Cadaver 8 | 72 | Male | White | 167.6 | 72.6 | 25.82 | Debility; Alzheimer's disease |
| Cadaver 9 | 79 | Male | White | 172.7 | 72.1 | 24.17 | Acute myocardial infarction; Coronary artery disease |
| Cadaver 10 | 80 | Male | White | 182.9 | 83.9 | 25.09 | Myocardial infarction; Cardiac arrest; Hypertension |
| Cadaver 11 | 86 | Female | White | 165.1 | 63.5 | 23.29 | Respiratory failure; Pneumonia |
| Cadaver 12 | 88 | Male | White | 177.8 | 68.0 | 21.52 | Natural causes; Unspecified |

**Table S3**: Kruskal-Wallis one-way ANOVA results, showing asymptotic significance (Asymp. Sig.) in chi-square with specified degrees of freedom (df) for the anterior (ACL) and posterior (PCL) cruciate ligaments, medial (MCL) and lateral (LCL) collateral ligaments material property data. ABREVIATIONS: $L_{0}$, original length of the ligament; $CSA$, cross-sectional area; $F$, force or load; $\sigma$, stress; $\varepsilon$, strain at the maximum point of the linear region (_max linear_) and failure (_failure_) point of the load-extension curve; $\boldsymbol{E}_{\boldsymbol{secant}}$, secant modulus; $\boldsymbol{E}_{\boldsymbol{tan}}$, tangent modulus; $k$, stiffness. *Significant at p≤0.05.

|  |  | $\boldsymbol{F}_{\boldsymbol{max linear}}$ **(N)** | $\boldsymbol{\sigma}_{\boldsymbol{max linear}}$ **(MPa)** | **Max Linear Extension (mm)** | $\boldsymbol{\varepsilon}_{\max\boldsymbol{linear}}$ **(%)** | $\boldsymbol{E}_{\boldsymbol{secant}}$ **(MPa)** | $\boldsymbol{E}_{\boldsymbol{tan}}$ **(MPa)** | $\boldsymbol{F}_{\boldsymbol{failure}}$ **(N)** | $\boldsymbol{\sigma}_{\boldsymbol{failure}}$  **(MPa)** | **Failure extension (mm)** | $\boldsymbol{\varepsilon}_{\boldsymbol{failure}}$ **(%)** | $\boldsymbol{k}$ **(N/mm)** |
| --- | --- | --- | --- | --- | --- | --- | --- | --- | --- | --- | --- | --- |
| **ACL** | **Chi-Square** | 6.846* | 3.923 | 7.615* | 4.027 | 3.154 | 2.577 | 3.808 | 3.769 | 6.179* | 3.103 | 3.513 |
|  | **df** | 2 | 2 | 2 | 2 | 2 | 2 | 2 | 2 | 2 | 2 | 2 |
|  | **Asymp. Sig.** | 0.033 | 0.141 | 0.022 | 0.134 | 0.207 | 0.276 | 0.149 | 0.152 | 0.046 | 0.212 | 0.173 |
| **PCL** | **Chi-Square** | 2.577 | 2.128 | 1.308 | 1.665 | 3.154 | 3.808 | 1.654 | 2.077 | 1.042 | 5.551 | 2.577 |
|  | **df** | 2 | 2 | 2 | 2 | 2 | 2 | 2 | 2 | 2 | 2 | 2 |
|  | **Asymp. Sig.** | 0.276 | 0.345 | 0.520 | 0.435 | 0.207 | 0.149 | 0.437 | 0.354 | 0.594 | 0.062 | 0.276 |
| **MCL** | **Chi-Square** | 2.76 | 2.373 | 0.893 | 4.093 | 2.16 | 3.133 | 0.573 | 2.333 | 0.454 | 0.84 | 0.773 |
|  | **df** | 2 | 2 | 2 | 2 | 2 | 2 | 2 | 2 | 2 | 2 | 2 |
|  | **Asymp. Sig.** | 0.252 | 0.305 | 0.640 | 0.129 | 0.340 | 0.209 | 0.751 | 0.311 | 0.797 | 0.657 | 0.679 |
| **LCL** | **Chi-Square** | 1.462 | 1.038 | 1.141 | 1.476 | 0.731 | 1.103 | 2.885 | 2.885 | 2.744 | 1.103 | 2.577 |
|  | **df** | 2 | 2 | 2 | 2 | 2 | 2 | 2 | 2 | 2 | 2 | 2 |
|  | **Asymp. Sig.** | 0.482 | 0.595 | 0.565 | 0.478 | 0.694 | 0.576 | 0.236 | 0.236 | 0.254 | 0.576 | 0.276 |

**Table S4**: Kendall’s Tau-b ($\tau_{b}$) correlation coefficient and their statistically significant (Sig.) values for the anterior (ACL) and posterior (PCL) cruciate ligaments, medial (MCL) and lateral (LCL) collateral ligaments material property data. ABREVIATIONS: N, specimen number; OA ICRS, osteoarthritis International Cartilage Repair Society; $L_{0}$, original length of the ligament; $CSA$, cross-sectional area; $F$, force or load; $\sigma$, stress; $\varepsilon$, strain at the maximum point of the linear region (_max linear_) and failure (_failure_) point of the load-extension curve; $\boldsymbol{E}_{\boldsymbol{secant}}$, secant modulus; $\boldsymbol{E}_{\boldsymbol{tan}}$, tangent modulus; $k$, stiffness. *Correlation is significant at the 0.05 level (2-tailed). **Correlation is significant at 0.01 level (2-tailed).

|  |  | | **Age** | **OA ICRS Grade** | $\boldsymbol{F}_{\boldsymbol{max linear}}$ **(N)** | $\boldsymbol{\sigma}_{\boldsymbol{max linear}}$ **(MPa)** | **Max Linear Extension (mm)** | $\boldsymbol{\varepsilon}_{\max\boldsymbol{linear}}$ **(%)** | $\boldsymbol{E}_{\boldsymbol{secant}}$ **(MPa)** | $\boldsymbol{E}_{\boldsymbol{tan}}$ **(MPa)** | $\boldsymbol{F}_{\boldsymbol{failure}}$ **(N)** | $\boldsymbol{\sigma}_{\boldsymbol{failure}}$  **(MPa)** | **Failure Extension (mm)** | $\boldsymbol{\varepsilon}_{\boldsymbol{failure}}$ **(%)** | $\boldsymbol{k}$ **(N/mm)** |
| --- | --- | --- | --- | --- | --- | --- | --- | --- | --- | --- | --- | --- | --- | --- | --- |
| **ACL** | **Age** | $\tau_{b}$ | 1.000 | .663** | -.626** | -.473* | -.443* | -0.277 | -0.412 | -0.382 | -.504* | -.534* | -.534* | -0.412 | -0.382 |
|  |  | Sig. (2-tailed) |  | 0.005 | 0.005 | 0.033 | 0.046 | 0.215 | 0.063 | 0.086 | 0.023 | 0.016 | 0.016 | 0.063 | 0.086 |
|  |  | N | 12 | 12 | 12 | 12 | 12 | 12 | 12 | 12 | 12 | 12 | 12 | 12 | 12 |
|  | **OA ICRS Grade** | $\tau_{b}$ | .663** | 1.000 | -.461* | -.526* | -.592* | -.497* | -0.395 | -0.362 | -0.197 | -0.362 | -0.263 | -0.132 | -0.197 |
|  |  | Sig. (2-tailed) | 0.005 |  | 0.048 | 0.024 | 0.011 | 0.034 | 0.090 | 0.120 | 0.397 | 0.120 | 0.259 | 0.572 | 0.397 |
|  |  | N | 12 | 12 | 12 | 12 | 12 | 12 | 12 | 12 | 12 | 12 | 12 | 12 | 12 |
| **PCL** | **Age** | $\tau_{b}$ | 1.000 | .663** | -0.412 | -0.382 | -0.229 | -0.140 | -.504* | -.534* | -0.321 | -0.382 | 0.031 | -0.076 | -0.412 |
|  |  | Sig. (2-tailed) |  | 0.005 | 0.063 | 0.086 | 0.303 | 0.534 | 0.023 | 0.016 | 0.149 | 0.086 | 0.890 | 0.731 | 0.063 |
|  |  | N | 12 | 12 | 12 | 12 | 12 | 12 | 12 | 12 | 12 | 12 | 12 | 12 | 12 |
|  | **OA ICRS Grade** | $\tau_{b}$ | .663** | 1.000 | -0.197 | -0.164 | -0.066 | -0.017 | -0.296 | -0.329 | -0.164 | -0.230 | 0.149 | -0.033 | -0.263 |
|  |  | Sig. (2-tailed) | 0.005 |  | 0.397 | 0.480 | 0.778 | 0.943 | 0.204 | 0.158 | 0.480 | 0.323 | 0.524 | 0.888 | 0.259 |
|  |  | N | 12 | 12 | 12 | 12 | 12 | 12 | 12 | 12 | 12 | 12 | 12 | 12 | 12 |
| **MCL** | **Age** | $\tau_{b}$ | 1.000 | .759** | -0.085 | -0.197 | 0.141 | 0.085 | -0.366 | -0.366 | -0.197 | -0.310 | -0.057 | -0.141 | -0.310 |
|  |  | Sig. (2-tailed) |  | 0.007 | 0.753 | 0.463 | 0.600 | 0.753 | 0.173 | 0.173 | 0.463 | 0.249 | 0.833 | 0.600 | 0.249 |
|  |  | N | 9 | 9 | 9 | 9 | 9 | 9 | 9 | 9 | 9 | 9 | 9 | 9 | 9 |
|  | **OA ICRS Grade** | $\tau_{b}$ | .759** | 1.000 | -0.210 | -0.269 | -0.150 | -0.210 | -0.269 | -0.269 | -0.090 | -0.210 | -0.121 | -0.269 | -0.090 |
|  |  | Sig. (2-tailed) | 0.007 |  | 0.451 | 0.333 | 0.590 | 0.451 | 0.333 | 0.333 | 0.747 | 0.451 | 0.665 | 0.333 | 0.747 |
|  |  | N | 9 | 9 | 9 | 9 | 9 | 9 | 9 | 9 | 9 | 9 | 9 | 9 | 9 |
| **LCL** | **Age** | $\tau_{b}$ | 1.000 | .663** | -0.321 | -0.290 | -0.137 | -0.308 | -0.137 | -0.168 | -0.412 | -0.198 | -0.015 | -0.168 | -.443* |
|  |  | Sig. (2-tailed) |  | 0.005 | 0.149 | 0.192 | 0.536 | 0.168 | 0.536 | 0.450 | 0.063 | 0.372 | 0.945 | 0.450 | 0.046 |
|  |  | N | 12 | 12 | 12 | 12 | 12 | 12 | 12 | 12 | 12 | 12 | 12 | 12 | 12 |
|  | **OA ICRS Grade** | $\tau_{b}$ | .663** | 1.000 | -0.164 | -0.263 | 0.066 | -0.182 | -0.263 | -0.296 | -0.296 | -.461* | -0.099 | -0.263 | -0.230 |
|  |  | Sig. (2-tailed) | 0.005 |  | 0.480 | 0.259 | 0.778 | 0.436 | 0.259 | 0.204 | 0.204 | 0.048 | 0.672 | 0.259 | 0.323 |
|  |  | N | 12 | 12 | 12 | 12 | 12 | 12 | 12 | 12 | 12 | 12 | 12 | 12 | 12 |

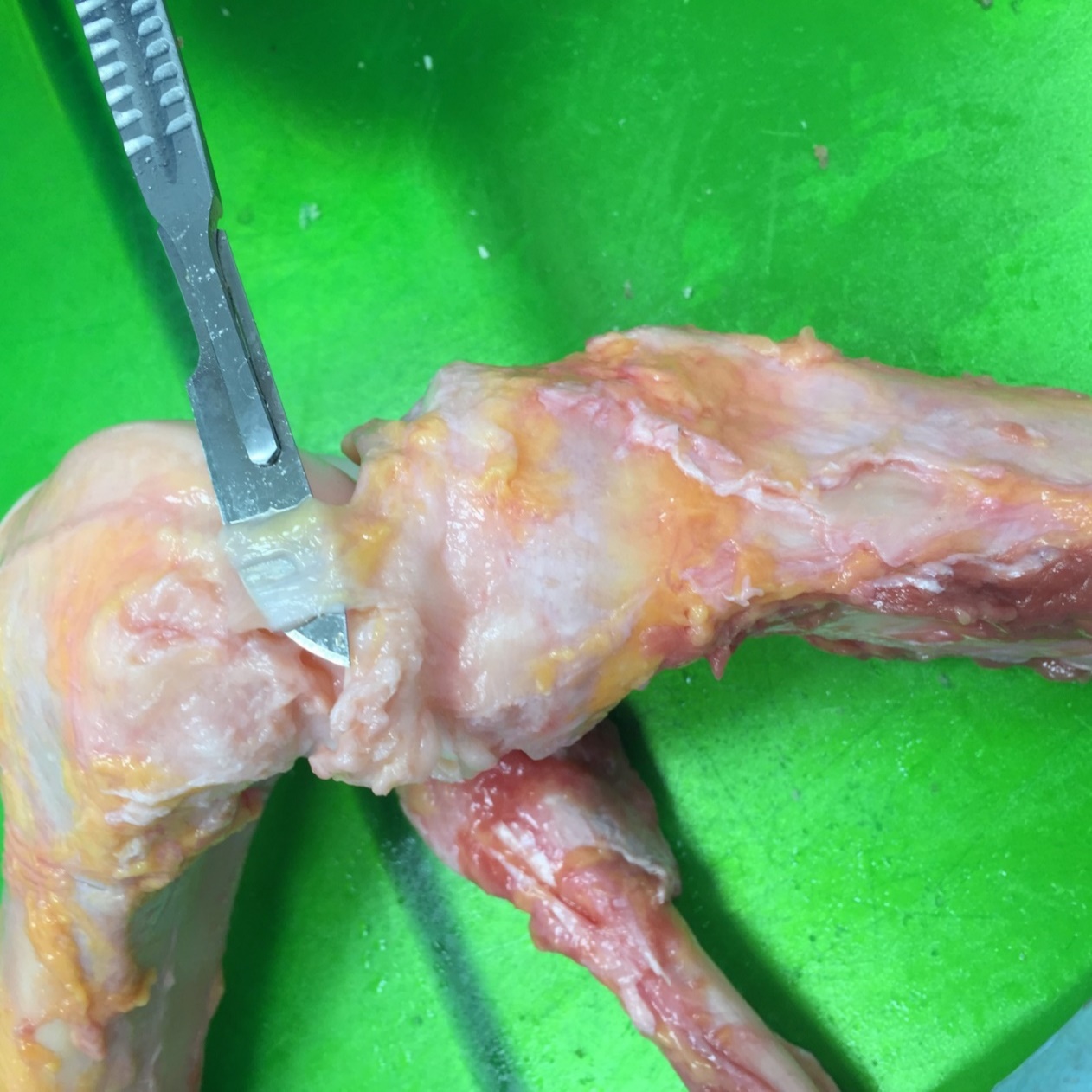

**Fig. S1.** A human medial collateral ligament from a young healthy donor that was severely abnormal, data from this ligament was excluded from statistical analyses.

| **ACL** |
| --- |
| **PCL** |
| **MCL** |
| **LCL** |
| **ACL** |
| **PCL** |
| **MCL** |
| **LCL** |
| **ACL** |
| **PCL** |
| **MCL** |
| **LCL** |
| **ACL** |
| **PCL** |
| **MCL** |
| **LCL** |
| **ACL** |
| **PCL** |
| **MCL** |
| **LCL** |

| Age (years) | Linear Force (N) |
| --- | --- |
| 31 | 399.336 |
| 37 | 777.074 |
| 43 | 337.071 |
| 49 | 169.611 |
| 51 | 267.291 |
| 58 | 112.842 |
| 72 | 300.102 |
| 72 | 229.336 |
| 79 | 72.03 |
| 80 | 100.938 |
| 86 | 82.7335 |
| 88 | 104.925 |

| Age (years) | Linear Stress (MPa) |
| --- | --- |
| 31 | 6.25894 |
| 37 | 16.0359 |
| 43 | 4.72923 |
| 49 | 4.8908 |
| 51 | 4.98315 |
| 58 | 1.17798 |
| 72 | 6.03207 |
| 72 | 2.25185 |
| 79 | 1.90679 |
| 80 | 1.3478 |
| 86 | 3.31172 |
| 88 | 1.63142 |

| Age (years) | Linear Force (N) |
| --- | --- |
| 31 | 446.364 |
| 37 | 765.825 |
| 43 | 270.116 |
| 49 | 630.721 |
| 51 | 300.83 |
| 58 | 750.58 |
| 72 | 327.815 |
| 72 | 873.644 |
| 79 | 260.954 |
| 80 | 49.5871 |
| 86 | 71.2168 |
| 88 | 182.972 |
|  | 527.406 |
|  | 219.739 |
|  | 294.365 |
|  | 303.288 |

| Age (years) | Linear Stress (MPa) |
| --- | --- |
| 31 | 5.15872 |
| 37 | 15.6853 |
| 43 | 3.52968 |
| 49 | 7.43849 |
| 51 | 5.07815 |
| 58 | 7.60716 |
| 72 | 5.24408 |
| 72 | 9.56511 |
| 79 | 3.71016 |
| 80 | 0.321477 |
| 86 | 1.07034 |
| 88 | 1.92089 |
|  | 7.41625 |
|  | 4.33779 |
|  | 3.63868 |
|  | 3.41339 |

| Age (years) | Linear Force (N) |
| --- | --- |
| 37 | 39.5582 |
| 43 | 288.398 |
| 51 | 235.566 |
| 58 | 66.615 |
| 72 | 188.087 |
| 72 | 96.5297 |
| 79 | 238.77 |
| 80 | 327.548 |
| 88 | 153.518 |
|  | 157.535 |
|  | 123.016 |
|  | 200.89 |
|  | 87.7157 |

| Age (years) | Linear Stress (MPa) |
| --- | --- |
| 37 | 2.17454 |
| 43 | 10.1962 |
| 51 | 5.7211 |
| 58 | 2.3127 |
| 72 | 5.62958 |
| 72 | 1.64491 |
| 79 | 6.0338 |
| 80 | 11.8604 |
| 88 | 4.31823 |
|  | 5.10113 |
|  | 3.772 |
|  | 5.89738 |
|  | 3.74916 |

| Age (years) | Linear Force (N) |
| --- | --- |
| 37 | 260.456 |
| 43 | 315.822 |
| 49 | 178.58 |
| 51 | 274.388 |
| 58 | 340.367 |
| 72 | 299.492 |
| 72 | 238.982 |
| 79 | 452.893 |
| 80 | 200.543 |
| 86 | 63.5942 |
| 88 | 80.8638 |
|  | 273.923 |
|  | 62.1334 |
|  | 222.728 |
|  | 144.965 |

| Age (years) | Linear Stress (MPa) |
| --- | --- |
| 37 | 3.63015 |
| 43 | 24.6188 |
| 49 | 4.47434 |
| 51 | 6.07759 |
| 58 | 9.44625 |
| 72 | 4.53288 |
| 72 | 5.37528 |
| 79 | 23.8572 |
| 80 | 4.01005 |
| 86 | 4.48858 |
| 88 | 3.26334 |
|  | 9.64943 |
|  | 8.65873 |
|  | 7.58789 |
|  | 8.00035 |

| Age (years) | Linear Strain (%) |
| --- | --- |
| 31 | 9.6 |
| 37 | 13.7 |
| 43 | 13.7 |
| 49 | 8.6 |
| 51 | 17.8 |
| 58 | 9.2 |
| 72 | 7.3 |
| 72 | 6.8 |
| 79 | 9.9 |
| 80 | 4.9 |
| 86 | 7.4 |
| 88 | 10.6 |

| Age (years) | Secant Modulus (MPa) |
| --- | --- |
| 31 | 65.1973 |
| 37 | 117.05 |
| 43 | 34.5199 |
| 49 | 56.8698 |
| 51 | 27.9952 |
| 58 | 12.8042 |
| 72 | 82.6311 |
| 72 | 33.1154 |
| 79 | 19.2605 |
| 80 | 27.5061 |
| 86 | 44.753 |
| 88 | 15.3908 |

| Age (years) | Linear Strain (%) |
| --- | --- |
| 31 | 9.1 |
| 37 | 17.4 |
| 43 | 17.3 |
| 49 | 11.2 |
| 51 | 11.9 |
| 58 | 11.9 |
| 72 | 15.2 |
| 72 | 13.8 |
| 79 | 12 |
| 80 | 7.3 |
| 86 | 7.4 |
| 88 | 13.8 |
|  | 13.1333 |
|  | 3.42384 |
|  | 11.5833 |
|  | 3.43303 |

| Age (years) | Secant Modulus (MPa) |
| --- | --- |
| 31 | 56.6893 |
| 37 | 90.1452 |
| 43 | 20.4028 |
| 49 | 66.4151 |
| 51 | 42.6736 |
| 58 | 63.9257 |
| 72 | 34.5005 |
| 72 | 69.3124 |
| 79 | 30.918 |
| 80 | 4.40379 |
| 86 | 14.4641 |
| 88 | 13.9195 |
|  | 56.7086 |
|  | 23.5771 |
|  | 27.9197 |
|  | 23.2194 |

| Age (years) | Linear Strain (%) |
| --- | --- |
| 37 | 9 |
| 43 | 7.5 |
| 51 | 5 |
| 58 | 2.3 |
| 72 | 4.7 |
| 72 | 5.2 |
| 79 | 6.3 |
| 80 | 5.3 |
| 88 | 5.8 |
|  | 5.95 |
|  | 2.93995 |
|  | 5.46 |
|  | 0.610737 |

| Age (years) | Secant Modulus (MPa) |
| --- | --- |
| 37 | 24.1616 |
| 43 | 135.949 |
| 51 | 114.422 |
| 58 | 100.552 |
| 72 | 119.778 |
| 72 | 31.6328 |
| 79 | 95.7746 |
| 80 | 223.781 |
| 88 | 74.4523 |
|  | 93.7712 |
|  | 48.6377 |
|  | 109.084 |
|  | 71.8416 |

| Age (years) | Linear Strain (%) |
| --- | --- |
| 37 | 11.2 |
| 43 | 14.3 |
| 49 | 11.4 |
| 51 | 11.9 |
| 58 | 8.7 |
| 72 | 15.4 |
| 72 | 8.4 |
| 79 | 11.7 |
| 80 | 8.7 |
| 86 | 5.4 |
| 88 | 8.5 |
|  | 11.5 |
|  | 1.99625 |
|  | 9.68333 |
|  | 3.43827 |

| Age (years) | Secant Modulus (MPa) |
| --- | --- |
| 37 | 32.4121 |
| 43 | 172.159 |
| 49 | 39.2486 |
| 51 | 51.0722 |
| 58 | 108.578 |
| 72 | 29.4343 |
| 72 | 63.9914 |
| 79 | 203.908 |
| 80 | 46.0925 |
| 86 | 83.1219 |
| 88 | 38.3922 |
|  | 80.694 |
|  | 59.3079 |
|  | 77.49 |
|  | 64.837 |

| Age (years) | Tangent Modulus (MPa) |
| --- | --- |
| 31 | 122.594 |
| 37 | 167.363 |
| 43 | 65.6335 |
| 49 | 144.99 |
| 51 | 36.6071 |
| 58 | 25.0968 |
| 72 | 138.908 |
| 72 | 56.7606 |
| 79 | 46.4689 |
| 80 | 41.7415 |
| 86 | 83.0776 |
| 88 | 30.6637 |

| Age (years) | Stiffness (N/mm) |
| --- | --- |
| 31 | 103.994 |
| 37 | 189.069 |
| 43 | 74.5567 |
| 49 | 49.3054 |
| 51 | 53.6299 |
| 58 | 29.9157 |
| 72 | 120.911 |
| 72 | 116.296 |
| 79 | 22.7367 |
| 80 | 54.2094 |
| 86 | 37.2673 |
| 88 | 29.9957 |

| Age (years) | Tangent Modulus (MPa) |
| --- | --- |
| 31 | 101.402 |
| 37 | 192.831 |
| 43 | 38.0818 |
| 49 | 124.918 |
| 51 | 73.9056 |
| 58 | 103.398 |
| 72 | 52.6019 |
| 72 | 113.399 |
| 79 | 48.7744 |
| 80 | 6.94979 |
| 86 | 37.0634 |
| 88 | 32.235 |
|  | 105.756 |
|  | 52.0923 |
|  | 48.5039 |
|  | 35.6385 |

| Age (years) | Stiffness (N/mm) |
| --- | --- |
| 31 | 136.253 |
| 37 | 146.71 |
| 43 | 52.0455 |
| 49 | 127.987 |
| 51 | 74.3524 |
| 58 | 137.117 |
| 72 | 52.6019 |
| 72 | 204.218 |
| 79 | 67.9567 |
| 80 | 19.4079 |
| 86 | 22.3811 |
| 88 | 38.9965 |
|  | 112.411 |
|  | 39.2181 |
|  | 67.5937 |
|  | 69.39 |

| Age (years) | Tangent Modulus (MPa) |
| --- | --- |
| 37 | 40.9132 |
| 43 | 283.424 |
| 51 | 181.679 |
| 58 | 179.751 |
| 72 | 255.184 |
| 72 | 55.0842 |
| 79 | 148.536 |
| 80 | 342.27 |
| 88 | 160.171 |
|  | 171.442 |
|  | 99.585 |
|  | 192.249 |
|  | 109.797 |

| Age (years) | Stiffness (N/mm) |
| --- | --- |
| 37 | 10.9884 |
| 43 | 35.6047 |
| 51 | 41.3274 |
| 58 | 22.8055 |
| 72 | 33.0732 |
| 72 | 37.1268 |
| 79 | 31.5833 |
| 80 | 53.2771 |
| 88 | 22.0571 |
|  | 27.6815 |
|  | 13.5576 |
|  | 35.4235 |
|  | 11.4082 |

| Age (years) | Tangent Modulus (MPa) |
| --- | --- |
| 37 | 51.9059 |
| 43 | 346.259 |
| 49 | 82.1694 |
| 51 | 88.9195 |
| 58 | 162.801 |
| 72 | 46.5037 |
| 72 | 102.912 |
| 79 | 294.787 |
| 80 | 87.1618 |
| 86 | 159.793 |
| 88 | 71.3552 |
|  | 146.411 |
|  | 118.924 |
|  | 127.086 |
|  | 90.5221 |

| Age (years) | Stiffness (N/mm) |
| --- | --- |
| 37 | 42.2819 |
| 43 | 36.2057 |
| 49 | 31.3298 |
| 51 | 49.0591 |
| 58 | 67.4529 |
| 72 | 32.4125 |
| 72 | 41.8386 |
| 79 | 62.4336 |
| 80 | 31.1498 |
| 86 | 19.6278 |
| 88 | 16.4024 |
|  | 45.2659 |
|  | 14.07 |
|  | 33.9775 |
|  | 16.714 |

| Age (years) | Failure Load (N) |
| --- | --- |
| 31 | 629.94 |
| 37 | 1554.16 |
| 43 | 503.059 |
| 49 | 280.963 |
| 51 | 309.096 |
| 58 | 205.601 |
| 72 | 808.337 |
| 72 | 613.771 |
| 79 | 161.699 |
| 80 | 373.937 |
| 86 | 84.766 |
| 88 | 144.403 |

| Age (years) | Failure Stress (MPa) |
| --- | --- |
| 31 | 9.87329 |
| 37 | 32.0544 |
| 43 | 7.10592 |
| 49 | 8.06937 |
| 51 | 5.74621 |
| 58 | 2.13713 |
| 72 | 16.2311 |
| 72 | 6.01745 |
| 79 | 2.79392 |
| 80 | 4.98499 |
| 86 | 3.3466 |
| 88 | 2.22379 |

| Age (years) | Failure Load (N) |
| --- | --- |
| 31 | 907.231 |
| 37 | 1336.49 |
| 43 | 369.202 |
| 49 | 923.9 |
| 51 | 378.242 |
| 58 | 1121.88 |
| 72 | 497.588 |
| 72 | 1432.7 |
| 79 | 601.741 |
| 80 | 148.143 |
| 86 | 227.51 |
| 88 | 274.387 |
|  | 839.49 |
|  | 393.036 |
|  | 530.345 |
|  | 474.073 |

| Age (years) | Failure Stress (MPa) |
| --- | --- |
| 31 | 10.4851 |
| 37 | 27.3486 |
| 43 | 4.83136 |
| 49 | 10.8872 |
| 51 | 6.37475 |
| 58 | 11.3604 |
| 72 | 7.94606 |
| 72 | 15.6772 |
| 79 | 8.53585 |
| 80 | 0.955036 |
| 86 | 3.4009 |
| 88 | 2.86673 |
|  | 11.8812 |
|  | 8.03081 |
|  | 6.56363 |
|  | 5.36627 |

| Age (years) | Failure Load (N) |
| --- | --- |
| 37 | 41.6043 |
| 43 | 387.705 |
| 51 | 280.806 |
| 58 | 514.051 |
| 72 | 458.466 |
| 72 | 139.019 |
| 79 | 326.724 |
| 80 | 399.005 |
| 88 | 285.142 |
|  | 306.042 |
|  | 200.417 |
|  | 321.671 |
|  | 121.876 |

| Age (years) | Failure Stress (MPa) |
| --- | --- |
| 37 | 2.23908 |
| 43 | 13.7426 |
| 51 | 6.78823 |
| 58 | 17.8132 |
| 72 | 13.6971 |
| 72 | 2.36895 |
| 79 | 8.25645 |
| 80 | 14.4091 |
| 88 | 8.11493 |
|  | 10.1458 |
|  | 6.96456 |
|  | 9.36929 |
|  | 4.89759 |

| Age (years) | Failure Load (N) |
| --- | --- |
| 37 | 479.726 |
| 43 | 426.516 |
| 49 | 279.4 |
| 51 | 341.772 |
| 58 | 629.387 |
| 72 | 371.209 |
| 72 | 121.309 |
| 79 | 591.602 |
| 80 | 299.436 |
| 86 | 229.028 |
| 88 | 169.349 |
|  | 431.36 |
|  | 134.81 |
|  | 296.989 |
|  | 169.756 |

| Age (years) | Failure Stress (MPa) |
| --- | --- |
| 37 | 6.67629 |
| 43 | 33.2108 |
| 49 | 6.97331 |
| 51 | 7.54733 |
| 58 | 17.4389 |
| 72 | 5.60089 |
| 72 | 2.70871 |
| 79 | 31.1223 |
| 80 | 5.96677 |
| 86 | 16.076 |
| 88 | 6.91354 |
|  | 14.3693 |
|  | 11.4548 |
|  | 11.398 |
|  | 10.6738 |

| Age (years) | Failure Strain (%) |
| --- | --- |
| 31 | 14.1665 |
| 37 | 27 |
| 43 | 21.869 |
| 49 | 11.9166 |
| 51 | 19.9997 |
| 58 | 13.699 |
| 72 | 16.2745 |
| 72 | 16.0343 |
| 79 | 12.3958 |
| 80 | 14.9558 |
| 86 | 7.44167 |
| 88 | 12.6446 |

| Age (years) | Length (mm) |
| --- | --- |
| 31 | 40 |
| 37 | 30 |
| 43 | 32 |
| 49 | 40 |
| 51 | 28 |
| 58 | 41 |
| 72 | 34 |
| 72 | 29 |
| 79 | 32 |
| 80 | 38 |
| 86 | 30 |
| 88 | 33 |

| Age (years) | Failure Strain (%) |
| --- | --- |
| 31 | 10.4851 |
| 37 | 27.3486 |
| 43 | 4.83136 |
| 49 | 10.8872 |
| 51 | 6.37475 |
| 58 | 11.3604 |
| 72 | 7.94606 |
| 72 | 15.6772 |
| 79 | 8.53585 |
| 80 | 0.955036 |
| 86 | 3.4009 |
| 88 | 2.86673 |
|  | 11.8812 |
|  | 8.03081 |
|  | 6.56363 |
|  | 5.36627 |

| Age (years) | Length (mm) |
| --- | --- |
| 31 | 36 |
| 37 | 30 |
| 43 | 30 |
| 49 | 44 |
| 51 | 34 |
| 58 | 46 |
| 72 | 41 |
| 72 | 31 |
| 79 | 32 |
| 80 | 35 |
| 86 | 43 |
| 88 | 34 |
|  | 36.6667 |
|  | 6.88961 |
|  | 36 |
|  | 4.89898 |

| Age (years) | Failure Strain (%) |
| --- | --- |
| 37 | 12.1669 |
| 43 | 9.73774 |
| 51 | 6.35959 |
| 58 | 9.65881 |
| 72 | 8.70536 |
| 72 | 7.63356 |
| 79 | 8.11151 |
| 80 | 6.14936 |
| 88 | 8.76402 |
|  | 9.48076 |
|  | 2.38431 |
|  | 7.87276 |
|  | 1.0691 |

| Age (years) | Length (mm) |
| --- | --- |
| 37 | 40 |
| 43 | 108 |
| 49 | 101 |
| 51 | 114 |
| 58 | 127 |
| 72 | 121 |
| 72 | 110 |
| 79 | 120 |
| 80 | 116 |
| 86 | 105 |
| 88 | 120 |
|  | 98 |
|  | 33.8009 |
|  | 115.333 |
|  | 6.50128 |

| Age (years) | Failure Strain (%) |
| --- | --- |
| 37 | 17.8183 |
| 43 | 17.3499 |
| 49 | 14.4 |
| 51 | 13.6878 |
| 58 | 15.2299 |
| 72 | 17.6645 |
| 72 | 5.6615 |
| 79 | 14.516 |
| 80 | 12.072 |
| 86 | 14.9445 |
| 88 | 14.1091 |
|  | 15.6972 |
|  | 1.81446 |
|  | 13.1613 |
|  | 4.08964 |

| Age (years) | Length (mm) |
| --- | --- |
| 37 | 55 |
| 43 | 61 |
| 49 | 52 |
| 51 | 47 |
| 58 | 58 |
| 72 | 60 |
| 72 | 68 |
| 79 | 62 |
| 80 | 74 |
| 86 | 60 |
| 88 | 58 |
|  | 54.6 |
|  | 5.41295 |
|  | 63.6667 |
|  | 6.121 |

**Fig. S2.** This figure shows the effect of age on material parameters of the anterior cruciate ligament (ACL), posterior cruciate ligament (PCL), medial collateral ligament (MCL) and lateral collateral ligament (LCL).

| **ACL** | 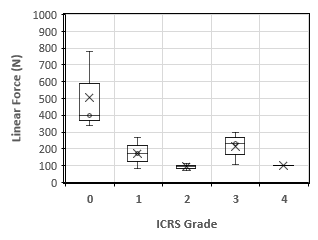 | 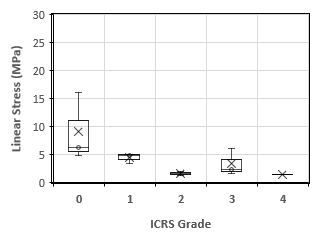 |
| --- | --- | --- |
| **PCL** | 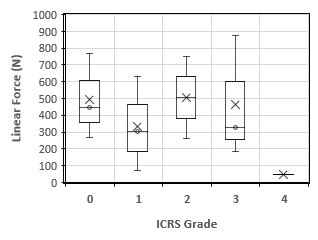 | 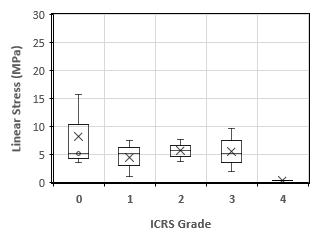 |
| **MCL** | 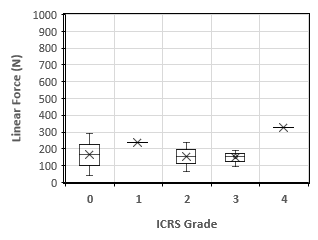 | 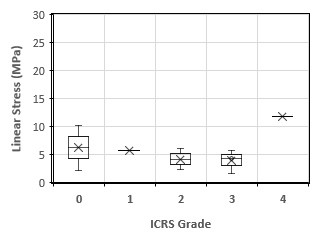 |
| **LCL** | 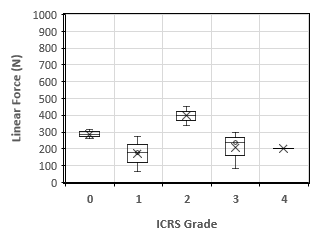 | 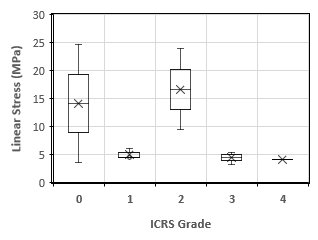 |

| **ACL** | 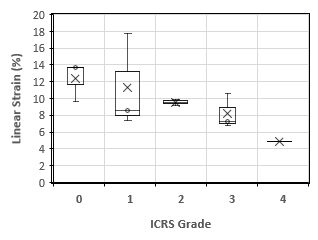 | 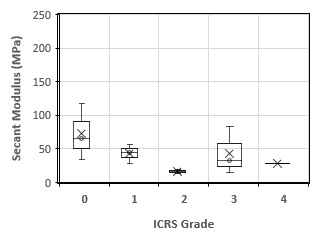 |
| --- | --- | --- |
| **PCL** | 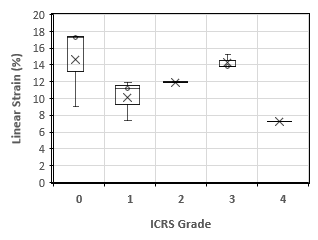 | 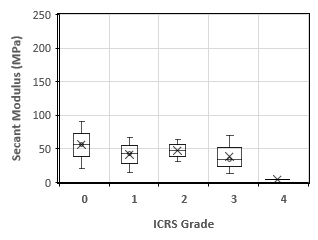 |
| **MCL** | 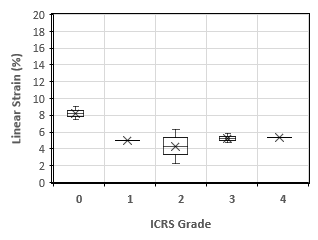 | 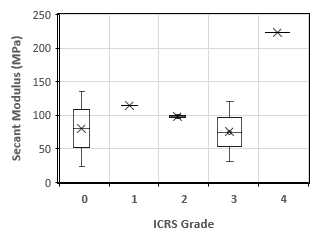 |
| **LCL** | 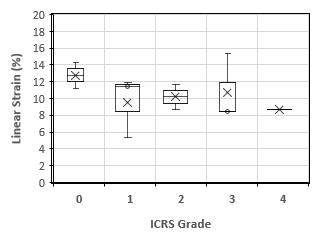 | 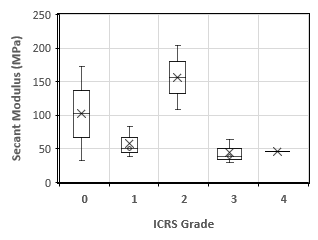 |
| **ACL** | 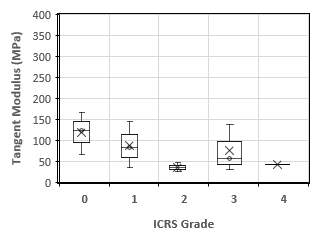 | 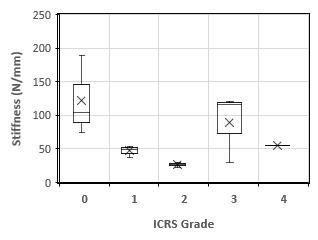 |
| **PCL** | 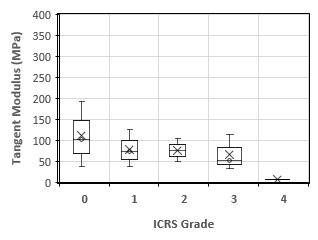 | 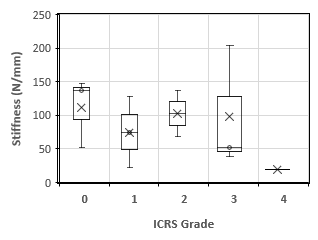 |
| **MCL** | 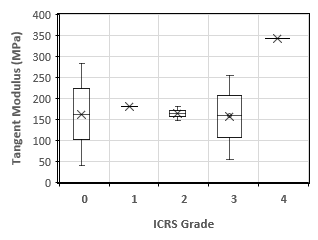 | 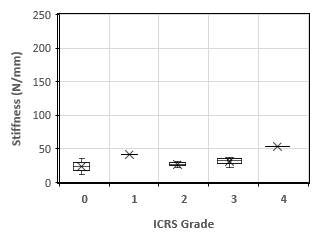 |
| **LCL** | 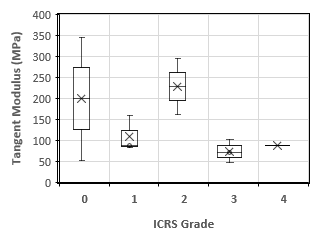 | 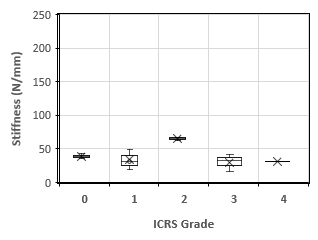 |

| **ACL** | 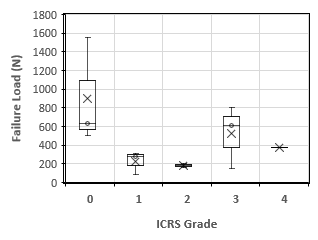 | 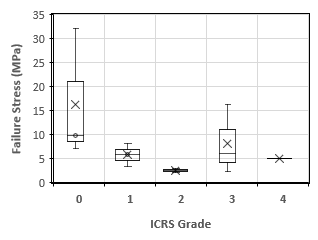 |
| --- | --- | --- |
| **PCL** | 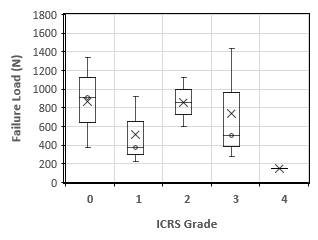 | 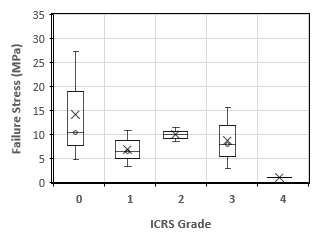 |
| **MCL** | 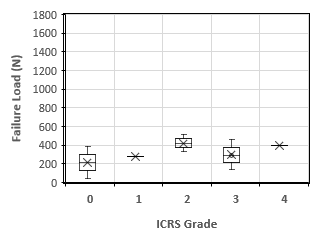 | 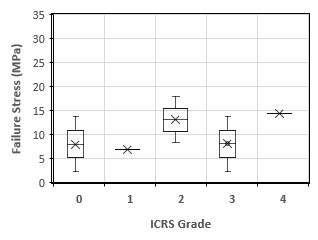 |
| **LCL** | 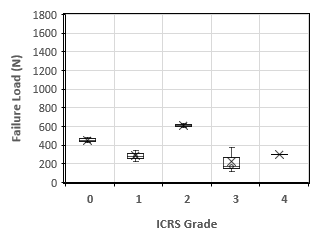 | 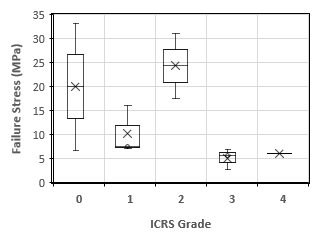 |

| **ACL** | 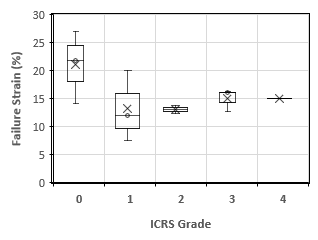 | 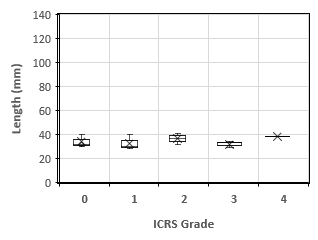 |
| --- | --- | --- |
| **PCL** | 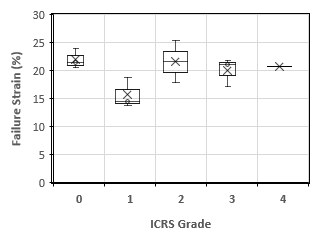 | 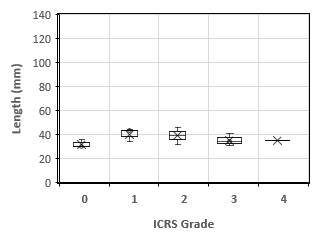 |
| **MCL** | 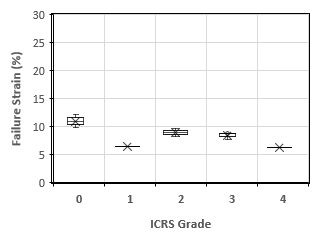 | 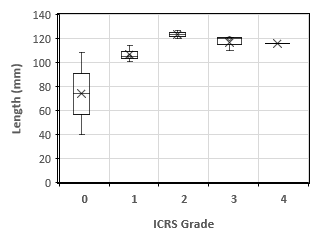 |
| **LCL** | 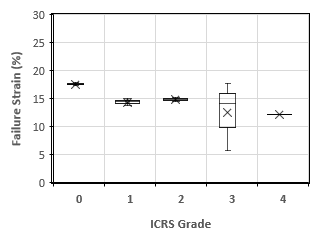 | 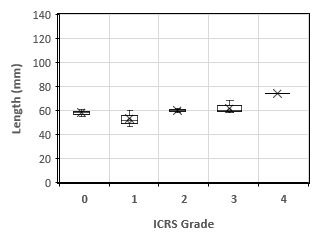 |

**Fig. S3.** This figure shows the effect of osteoarthritis (International Cartilage Repair Society (ICRS) Grading 0 to 4) on material parameters of the anterior cruciate ligament (ACL), posterior cruciate ligament (PCL), medial collateral ligament (MCL) and lateral collateral ligament (LCL).

| **Healthy knees (ICRS grade = 0)** | **OA knees (ICRS grades 1 to 4)** |
| --- | --- |

Linear Stress (MPa)

| Age (Years) | ACL | PCL |
| --- | --- | --- |
| 31 | 6.25894 | 5.15872 |
| 37 | 16.0359 | 15.6853 |
| 43 | 4.72923 | 3.52968 |
| 86 | 3.31172 | 1.07034 |

| Age (Years) | MCL | LCL |
| --- | --- | --- |
| 37 | 2.17454 | 3.63015 |
| 43 | 10.1962 | 24.6188 |
| 86 |  | 4.48858 |

Linear Stress (MPa)

| Age (Years) | ACL | PCL | MCL | LCL |
| --- | --- | --- | --- | --- |
| 49 | 4.8908 | 7.43849 |  | 4.47434 |
| 51 | 4.98315 | 5.07815 | 5.7211 | 6.07759 |
| 58 | 1.17798 | 7.60716 | 2.3127 | 9.44625 |
| 72 | 6.03207 | 5.24408 | 5.62958 | 4.53288 |
| 72 | 2.25185 | 9.56511 | 1.64491 | 5.37528 |
| 79 | 1.90679 | 3.71016 | 6.0338 | 23.8572 |
| 80 | 1.3478 | 0.321477 | 11.8604 | 4.01005 |
| 88 | 1.63142 | 1.92089 | 4.31823 | 3.26334 |

Linear Strain (%)

| Age (Years) | ACL | PCL |
| --- | --- | --- |
| 31 | 9.6 | 9.1 |
| 37 | 13.7 | 17.4 |
| 43 | 13.7 | 17.3 |
| 86 | 7.4 | 7.4 |

| Age (Years) | MCL | LCL |
| --- | --- | --- |
| 37 | 9 | 11.2 |
| 43 | 7.5 | 14.3 |
| 86 |  | 5.4 |

Linear Strain (%)

| Age (Years) | ACL | PCL | MCL | LCL |
| --- | --- | --- | --- | --- |
| 49 | 8.6 | 11.2 |  | 11.4 |
| 51 | 17.8 | 11.9 | 5 | 11.9 |
| 58 | 9.2 | 11.9 | 2.3 | 8.7 |
| 72 | 7.3 | 15.2 | 4.7 | 15.4 |
| 72 | 6.8 | 13.8 | 5.2 | 8.4 |
| 79 | 9.9 | 12 | 6.3 | 11.7 |
| 80 | 4.9 | 7.3 | 5.3 | 8.7 |
| 88 | 10.6 | 13.8 | 5.8 | 8.5 |

| **Healthy knees (ICRS grade = 0)** | **OA knees (ICRS grades 1 to 4)** |
| --- | --- |

| **Healthy knees (ICRS grade = 0)** | **OA knees (ICRS grades 1 to 4)** |
| --- | --- |

**Fig. S4.** Trends showing the effect of ageing in healthy and osteoarthritis (OA) cadaveric knees on the tensile properties of the anterior cruciate ligament (ACL), posterior cruciate ligament (PCL), medial collateral ligament (MCL) and lateral collateral ligament (LCL).

(A) **Young healthy cadaver**
(B) **Old OA cadaver**

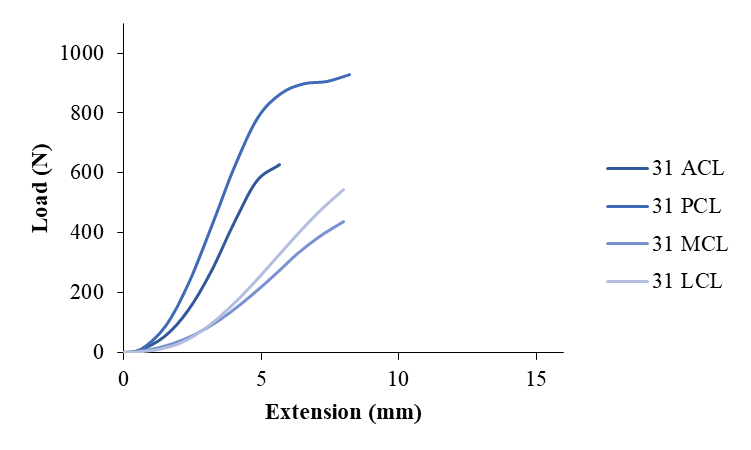

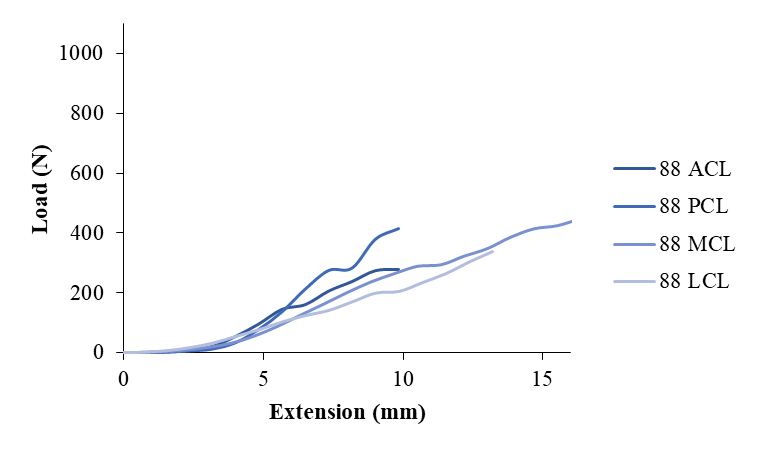

Load (N)

| Extension (mm) | 31 ACL |
| --- | --- |
| 0 | 0 |
| -0.00011 | 0.00582 |
| -0.00017 | -0.00695 |
| -0.00017 | -0.02626 |
| -0.00017 | -0.02195 |
| 0.60067 | 8.63802 |
| 1.49362 | 54.0492 |
| 2.33246 | 139.994 |
| 3.16636 | 268.724 |
| 3.99982 | 431.249 |
| 4.8331 | 573.776 |
| 5.66644 | 628.94 |

| Extension (mm) | 31 PCL |
| --- | --- |
| 0 | 0 |
| 6e-05 | 0.06711 |
| 6e-05 | 0.08554 |
| 6e-05 | 0.05203 |
| 6e-05 | 0.05211 |
| 0.64045 | 12.276 |
| 1.52762 | 89.7984 |
| 2.36606 | 233.867 |
| 3.19986 | 424.52 |
| 4.03331 | 623.484 |
| 4.86659 | 785.105 |
| 5.69999 | 865.341 |
| 6.53333 | 898.512 |
| 7.36673 | 906.052 |
| 8.20001 | 929.087 |

| Extension (mm) | 31 MCL |
| --- | --- |
| 0 | 0 |
| 0 | -0.08738 |
| 0 | -0.09561 |
| -5e-05 | 0.00522 |
| -5e-05 | 0.00337 |
| 0.39219 | 1.50518 |
| 1.32345 | 16.6335 |
| 2.16563 | 42.1457 |
| 2.99988 | 79.9722 |
| 3.83328 | 130.664 |
| 4.66662 | 192.325 |
| 5.4999 | 259.719 |
| 6.33318 | 331.351 |
| 7.16652 | 390.302 |
| 7.99998 | 437.484 |

| Extension (mm) | 31 LCL |
| --- | --- |
| 0 | 0 |
| -0.00012 | -0.00719 |
| 0 | 0.07013 |
| 0 | 0.00636 |
| 0 | 0.011 |
| 0.39207 | 1.08207 |
| 1.32379 | 11.0912 |
| 2.1658 | 35.4948 |
| 2.99987 | 82.5966 |
| 3.83333 | 147.966 |
| 4.6665 | 225.587 |
| 5.49989 | 310.882 |
| 6.33329 | 396.721 |
| 7.16663 | 476.556 |
| 7.99991 | 544.821 |

Load (N)

| Extension (mm) | 88 ACL |
| --- | --- |
| 0 | 0 |
| 0.00499 | 0.02239 |
| 0.0051 | 0.01571 |
| 0.60815 | 0.44585 |
| 1.49991 | 1.76233 |
| 2.33869 | 7.20821 |
| 3.1726 | 20.5798 |
| 4.00605 | 54.1524 |
| 4.83928 | 95.8146 |
| 5.67262 | 144.785 |
| 6.50596 | 160.601 |
| 7.3393 | 205.762 |
| 8.17264 | 237.29 |
| 9.00598 | 273.725 |
| 9.83938 | 277.856 |

| Extension (mm) | 88 PCL |
| --- | --- |
| 0 | 0 |
| -0.00029 | 0.01563 |
| -0.00029 | 0.00988 |
| 0.6197 | 0.54669 |
| 1.51045 | 2.01841 |
| 2.34906 | 5.82532 |
| 3.18296 | 12.0118 |
| 4.0163 | 33.5618 |
| 4.84964 | 78.1337 |
| 5.68298 | 137.295 |
| 6.51627 | 213.378 |
| 7.34961 | 275.084 |
| 8.183 | 282.914 |
| 9.01623 | 379.192 |
| 9.84957 | 414.766 |

| Extension (mm) | 88 MCL |
| --- | --- |
| 0 | 0 |
| 0 | 0.03324 |
| 0 | 0.03085 |
| 0.41338 | 0.5872 |
| 1.3405 | 3.22266 |
| 2.18246 | 8.19073 |
| 3.01659 | 17.3505 |
| 3.84999 | 31.6835 |
| 4.68338 | 55.6806 |
| 5.51667 | 88.2483 |
| 6.35012 | 126.149 |
| 7.1834 | 163.262 |
| 8.01674 | 201.538 |
| 8.84997 | 236.104 |
| 9.68337 | 263.648 |
| 10.5168 | 288.525 |
| 11.35 | 293.36 |
| 12.1834 | 322.39 |
| 13.0167 | 347.646 |
| 13.8499 | 385.7 |
| 14.6834 | 413.58 |
| 15.5167 | 424.083 |
| 16.35 | 447.562 |

| Extension (mm) | 88 LCL |
| --- | --- |
| 0 | 0 |
| 0 | 0.01345 |
| 0 | 0.00307 |
| 0 | 0.00561 |
| 0.62061 | 1.1837 |
| 1.5109 | 6.39167 |
| 2.3494 | 16.5985 |
| 3.18331 | 31.7921 |
| 4.0167 | 54.7399 |
| 4.85004 | 77.449 |
| 5.68327 | 103.941 |
| 6.51667 | 125.176 |
| 7.35001 | 142.614 |
| 8.18329 | 171.317 |
| 9.01669 | 200.373 |
| 9.85003 | 205.899 |
| 10.6832 | 235.68 |
| 11.5167 | 265.671 |
| 12.3499 | 304.954 |
| 13.1833 | 339.508 |

**Fig. S5.** Example of load (N) against extension (mm) in the anterior cruciate ligament (ACL), posterior cruciate ligament (PCL), medial collateral ligament (MCL) and lateral collateral ligament (LCL) in (A) young healthy cadaver (31 years, grade 0), (B) old osteoarthritis cadaver (88 years, grade 3).

(A) **Young healthy cadaver**
(B) **Old OA cadaver**

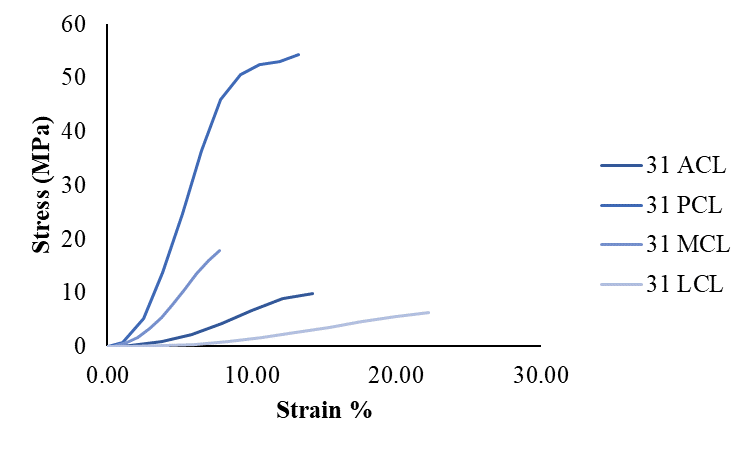

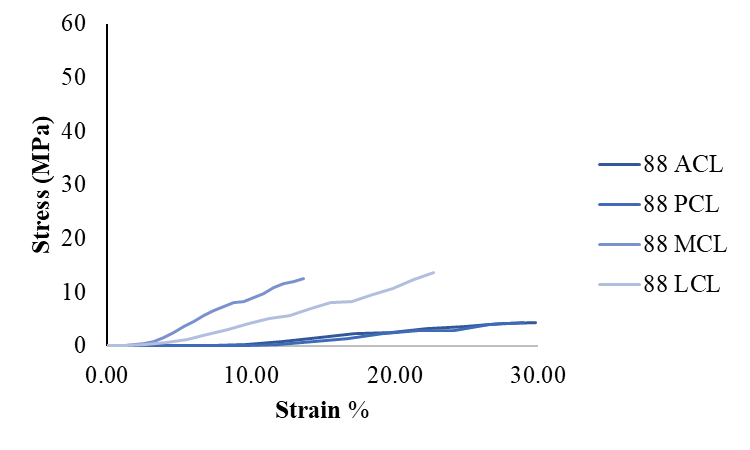

Stress (MPa)

| Strain % | 31 ACL |
| --- | --- |
| -0.000425 | -0.00034403 |
| 1.50168 | 0.135387 |
| 3.73405 | 0.847133 |
| 5.83115 | 2.19417 |
| 7.9159 | 4.21181 |
| 9.99955 | 6.75912 |
| 12.0828 | 8.993 |
| 14.1661 | 9.8576 |

| Strain % | 31 PCL |
| --- | --- |
| 9.67742e-05 | 0.00304683 |
| 1.03298 | 0.717771 |
| 2.4639 | 5.25045 |
| 3.81623 | 13.674 |
| 5.16106 | 24.8214 |
| 6.50534 | 36.4547 |
| 7.84934 | 45.9045 |
| 9.19353 | 50.5958 |
| 10.5376 | 52.5353 |
| 11.8818 | 52.9762 |
| 13.2258 | 54.323 |

| Strain % | 31 MCL |
| --- | --- |
| -4.85437e-05 | 0.000137498 |
| 0.380767 | 0.0614121 |
| 1.2849 | 0.678656 |
| 2.10255 | 1.71957 |
| 2.9125 | 3.26291 |
| 3.72163 | 5.33117 |
| 4.5307 | 7.84695 |
| 5.33971 | 10.5967 |
| 6.14872 | 13.5193 |
| 6.95779 | 15.9245 |
| 7.76697 | 17.8496 |

| Strain % | 31 LCL |
| --- | --- |
| 0 | 0.000127129 |
| 1.08908 | 0.0125057 |
| 3.67719 | 0.128184 |
| 6.01611 | 0.410222 |
| 8.33297 | 0.954587 |
| 10.6481 | 1.71008 |
| 12.9625 | 2.60715 |
| 15.2775 | 3.59294 |
| 17.5925 | 4.58499 |
| 19.9073 | 5.50767 |
| 22.222 | 6.29662 |

Stress (MPa)

| Strain % | 88 ACL |
| --- | --- |
| 0 | 0 |
| 0.0151212 | 0.00034813 |
| 0.0154545 | 0.000244267 |
| 1.84288 | 0.00693229 |
| 4.54518 | 0.0274015 |
| 7.08694 | 0.112077 |
| 9.61394 | 0.319985 |
| 12.1395 | 0.841987 |
| 14.6645 | 1.48977 |
| 17.1898 | 2.25119 |
| 19.715 | 2.4971 |
| 22.2403 | 3.19929 |
| 24.7656 | 3.68949 |
| 27.2908 | 4.25601 |
| 29.8163 | 4.32024 |

| Strain % | 88 PCL |
| --- | --- |
| 0 | 0 |
| -0.000852941 | 0.000164088 |
| -0.000852941 | 0.000103723 |
| 1.82265 | 0.00573932 |
| 4.4425 | 0.0211899 |
| 6.909 | 0.061156 |
| 9.36165 | 0.126104 |
| 11.8126 | 0.352342 |
| 14.2636 | 0.820271 |
| 16.7146 | 1.44136 |
| 19.1655 | 2.2401 |
| 21.6165 | 2.88792 |
| 24.0676 | 2.97012 |
| 26.5183 | 3.98088 |
| 28.9693 | 4.35434 |

| Strain % | 88 MCL |
| --- | --- |
| 0 | 0 |
| 0 | 0.000934995 |
| 0 | 0.000867767 |
| 0.344483 | 0.0165171 |
| 1.11708 | 0.0906489 |
| 1.81872 | 0.230394 |
| 2.51383 | 0.488044 |
| 3.20832 | 0.891213 |
| 3.90282 | 1.56622 |
| 4.59723 | 2.4823 |
| 5.29177 | 3.5484 |
| 5.98617 | 4.59234 |
| 6.68062 | 5.66899 |
| 7.37497 | 6.64129 |
| 8.06948 | 7.41606 |
| 8.76402 | 8.11579 |
| 9.45832 | 8.25182 |
| 10.1528 | 9.06837 |
| 10.8472 | 9.77878 |
| 11.5416 | 10.8492 |
| 12.2362 | 11.6334 |
| 12.9306 | 11.9289 |
| 13.625 | 12.5893 |

| Strain % | 88 LCL |
| --- | --- |
| 0 | 0 |
| 0 | 0.000542787 |
| 0 | 0.000123893 |
| 0 | 0.000226397 |
| 1.07002 | 0.0477693 |
| 2.605 | 0.257942 |
| 4.05069 | 0.669849 |
| 5.48847 | 1.283 |
| 6.92534 | 2.20908 |
| 8.36214 | 3.12553 |
| 9.79874 | 4.19463 |
| 11.2356 | 5.05159 |
| 12.6724 | 5.7553 |
| 14.1091 | 6.91366 |
| 15.546 | 8.08624 |
| 16.9828 | 8.30925 |
| 18.4194 | 9.5111 |
| 19.8563 | 10.7214 |
| 21.293 | 12.3067 |
| 22.7299 | 13.7012 |

**Fig. S6.** Example of stress (MPa) against strain (%) in the anterior cruciate ligament (ACL), posterior cruciate ligament (PCL), medial collateral ligament (MCL) and lateral collateral ligament (LCL) in (A) young healthy cadaver (31 years, grade 0), (B) old osteoarthritis cadaver (88 years, grade 3).

(A) **Young vs old**
(B) **Healthy vs OA**

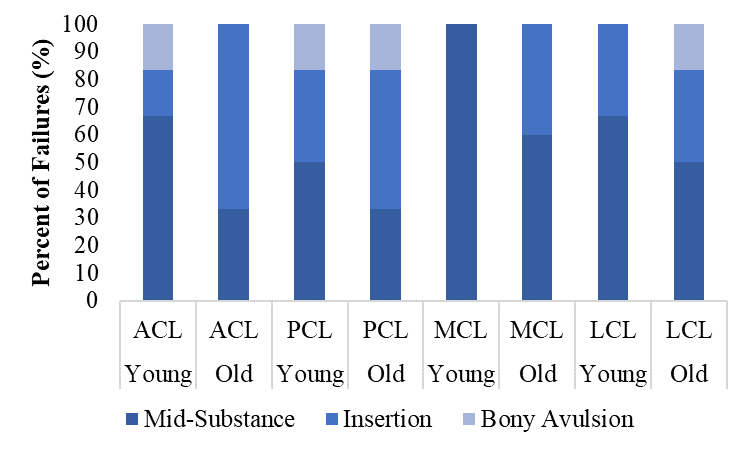

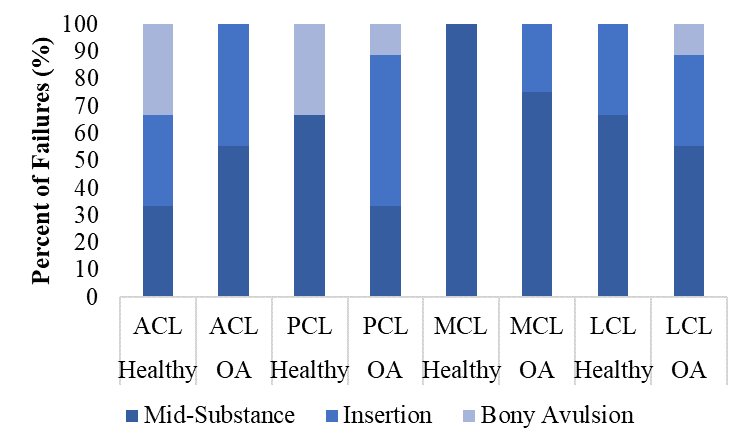

Percent of Failures (%)

|  | Mid-Substance | Insertion | Bony Avulsion |
| --- | --- | --- | --- |
| Young ACL | 66.6667 | 16.6667 | 16.6667 |
| Old ACL | 33.3333 | 66.6667 | 0 |
| Young PCL | 50 | 33.3333 | 16.6667 |
| Old PCL | 33.3333 | 50 | 16.6667 |
| Young MCL | 100 | 0 | 0 |
| Old MCL | 60 | 40 | 0 |
| Young LCL | 66.6667 | 33.3333 | 0 |
| Old LCL | 50 | 33.3333 | 16.6667 |

Percent of Failures (%)

|  | Mid-Substance | Insertion | Bony Avulsion |
| --- | --- | --- | --- |
| Healthy ACL | 33.3333 | 33.3333 | 33.3333 |
| OA ACL | 55.5556 | 44.4444 | 0 |
| Healthy PCL | 66.6667 | 0 | 33.3333 |
| OA PCL | 33.3333 | 55.5556 | 11.1111 |
| Healthy MCL | 100 | 0 | 0 |
| OA MCL | 75 | 25 | 0 |
| Healthy LCL | 66.6667 | 33.3333 | 0 |
| OA LCL | 55.5556 | 33.3333 | 11.1111 |

**Fig. S7.** (A) Failure site in percentage of young (31-58 years) versus old (72-88 years) cadavers for anterior cruciate ligament (ACL), posterior cruciate ligament (PCL), medial collateral ligament (MCL) and lateral collateral ligament LCL). (B) Failure site in percentage of healthy (grade 0) versus osteoarthritis (grade 1 – 4) for the ACL, PCL, MCL and LCL.
